# Perivascular matrix densification dysregulates angiogenesis and activates pro-inflammatory endothelial cells

**DOI:** 10.1101/2025.05.20.655215

**Authors:** Jingyi Xia, William Y. Wang, Kyle A. Jacobs, Daphne Lin, Evan H. Jarman, Daniel L. Matera, Kristen Loesel, Christopher D. Davidson, Harrison L. Hiraki, Xiaotian Tan, Eve H. Shikanov, Robert N. Kent, Carole Parent, Xudong Fan, Ariella Shikanov, Matthew L. Kutys, Brendon M. Baker

## Abstract

Fibrosis is central to numerous fatal conditions including solid cancers, pulmonary fibrosis, cirrhosis and post-infarct cardiac fibrosis amongst many others, thereby collectively contributing to 45% of all deaths in developed nations. The potential for fibrosis across most organ systems may stem from its connections to wound healing and the ubiquitous presence of vascular endothelium. Endothelial cells (ECs) and angiogenesis, cells and associated biological program central to wound healing, have been heavily implicated in many organ-specific fibroses, but the relationship between angiogenesis and fibrogenesis remains debated and little has been established in terms of how the EC phenotype governs tissue healing vs. fibrosis. Here, we examine a murine lung injury model enabling EC lineage tracing and observe the invasion of aberrant ECs from the bronchial microvasculature following lung injury along with concurrent densification of matrix fibers surrounding these vessels. To investigate the underlying mechanisms governing their appearance, we established a microphysiological system (MPS) of arteriole/venule-scale microvessels embedded within a tunable stromal mimetic matrix and find that heightened extracellular matrix fiber density activates ECs, drives endothelial to mesenchymal transition, and promotes aberrant tip EC (ATEC) invasion into the matrix. ATECs remain adherent to fibrotic matrix and possess a pro-inflammatory phenotype that secretes TGF-β2. Notably, our studies establish that the formation of ATECs is gated by destabilization of endothelial adherens junction upon EC adhesion to fibrous matrix, and associated regulation of TGF-β signaling that is mediated by a novel VE-cadherin – TGF-βR2 axis. The current lack of effective anti-fibrotic therapies suggests potential critical involvement of other cell types such as ECs, and our findings suggest new contributions of ECs to fibrotic progression that may better inform future targets for novel anti-fibrotic therapeutics.

## INTRODUCTION

Fibrotic tissue remodeling across sundry tissues and organ systems is often typified as a persistent or overexuberant wound healing response. While myofibroblasts (MF) have been regarded as the direct cellular driver of fibrotic matrix remodeling and therefore primary focus of therapeutic targeting to date, accumulating evidence has highlighted the role of other cell types including endothelial cells (ECs), epithelial cells, and immune cells including macrophages as additional contributors ^1^. ECs and the angiogenic process is central process to wound healing; as such, it is plausible that changes to the extracellular matrix (ECM) in a fibrotic microenvironment may derail the EC angiogenic response from acting to restore tissue homeostasis to instead promoting disease-driving cell function and associated signaling.

Supporting the involvement of ECs in fibrosis, histological examination of various murine models of fibrosis has consistently revealed microvascular abnormalities, but the direction (increased vs. decreased vascularity)^2–4^ underlying mechanisms, and the role of these changes in driving fibrosis not been established. *In vivo* lineage tracing in various murine models of fibrotic disease (eg. cardiac, cancer, lung, liver, and kidney) suggest as many as 30% of myofibroblasts originate from EC precursors that undergo endothelial to mesenchymal transition (EndMT) prior to myofibrogenesis^5–11^. Additionally, angiocrine signling from ECs have been shown to control a switch between tissue regeneration vs. fibrotic progression in acute injury models of lung and liver fibrosis^12–14^. Fibrosis consistently involves progressive increases in matrix density and crosslinking which alter or impair parenchymal cell function, ultimately culminating in organ failure. Still, it remains unclear if and how these biophysical changes to the ECM influence EC signaling and function after tissue injury and in fibrotic diseases.

Observations consistent across in vivo and in vitro models of angiogenesis support a critical, initiating step of EC activation and tip cell invasion into the surrounding interstitial matrix. Given evident parallels to epithelial cell invasion enabled by epithelial-mesenchymal transition, previous work has implicated endothelial-mesenchymal transition (EndMT) in the initiation of angiogenesis^15–19^. Interestingly, a separate body of work suggests ECs can directly contribute to fibrosis in vitro and in multiple organ systems via EndMT and subsequent myofibrogenesis ^10,11,20–27^. The established central role of MFs along with evidence that EndMT induces a multipotent EC phenotype has led many to explore whether ECs drive fibrotic progression by transdifferentiating into *bona fide* MFs. However, the varied identification of these cells in different organ fibroses despite consistent vascular abnormalities begs the question of whether ECs may drive fibrosis through alternative, yet equally important, mechanisms.

Previous work on EndMT has largely focused on biochemical and genetic mediators ^10,11,20–24^, however studies examining epithelial-mesenchymal transition indicate that physical ECM cues can potently modulate the impact of such signals ^28–31^. A parallel but currently unexplored concept likely extends to EndMT and the angiogenic response, which is supported by observations that angiogenesis is highly sensitive to physical properties of the surrounding ECM^32–35^. Interestingly, critical to angiogenesis is the dynamic regulation of vascular adherens junctions (AJs), which mechanically stabilize EC-EC connections and critically regulate effector signaling pathways governing EC fate and behavior^36,37^. Further, biochemical and mechanical cues mediate signaling at EC-ECM adhesions which in turn influence the assembly and stability of EC AJs. Thus, dissecting the putative relationship between physical and soluble microenvironmental cues that may underlie EC fate and function requires integrative experimental systems that can suitably approximate the native tissue microenvironment while providing experimental control.

Informed by observations in the murine bleomycin lung injury model, here we integrated tunable fibrous hydrogel composites and microphysiological systems to explore the influence of fibrotic matrix (ie. heightened perivascular fiber density) on arteriole/venule-scale endothelium and investigated EC-ECM interactions in the context of activating quiescent ECs from microvessels into invasive aberrant tip ECs (ATECs), which in other matrix settings typically lead angiogenic sprouts to extend capillaries. We demonstrate heightened fiber density via EC mechanosensing destabilizes EC AJs, diminishes vascular barrier function, and concomitantly increases ATEC formation via EndMT. Using unbiased proteomics, we identify fiber-induced AJ destabilization decreases a VE-cadherin – TGFβ-R2 interaction which underlies fiber-mediated enhancement of EC TGF-β signaling. Further, transcriptomic and secretomic characterization of ATECs revealed that these cells transition towards a secretory, pro-inflammatory phenotype, and are additionally a significant source of TGF-β2, a key pro-fibrotic growth factor that may act on macrophages, fibroblasts, but also ECs resident to parental vasculature. Indeed^10,22^, exposure of parent vessel endothelium to TGF-β2 leads to heightened EC apoptosis as a function of perivascular fiber density, consistent with observations of arteriolar/venular EC apoptosis in the bleomycin mouse model. Together, our studies delineate how changes in perivascular ECM fiber density destabilize EC AJs, promote EndMT, and the formation of inflammatory and pro-fibrotic ATECs. Furthermore, this work provides evidence for a fibrotic feedforward cycle mediated purely by ECs that may act in parallel or synergistically with myofibroblast driven fibrotic activity.

## RESULTS & DISCUSSION

### Characterization of EC expansion in a murine model of pulmonary fibrosis

To track the location and morphology of pulmonary ECs during injury-induced lung fibrogenesis, we developed a 3D lung tissue imaging pipeline combining a genetically engineered mouse model enabling EC lineage tracing, precision cut lung slices (PCLS, 200 μm thick), optical clearing methods, and volumetric confocal imaging. Adult Tie2-Cre/mTmG, where Tie2 (TEK)-driven Cre recombinase activity drives a permanent switch from membrane-localized tdTomato to GFP expression, were administered a single intratracheal instillation of bleomycin (0.04 U/mouse) to initiate fibrotic remodeling (ie. matrix deposition and crosslinking) which typically peaks in severity after 2-3 weeks. (**Figure 1a**). Following cardiac perfusion and lung extraction, PCLS were sectioned by vibratome and cleared using the CUBIC method^38^, immunostained for α-smooth muscle actin (αSMA), counterstained with DAPI and Alexa Fluor succinimidyl ester (AFSE, which fluorescently labels all amine-containing ECM and cellular proteins), and imaged by laser-scanning confocal microscopy (**Figure 1b**). Low magnification tile-scan imaging of entire lung sections across healthy control (day 0) and tissues at the peak of the fibrotic response (day 21) revealed marked changes in GFP^+^ EC and αSMA^+^ MF density resulting from bleomycin-induced lung injury. In control lungs, EC density within alveolar regions was uniform and αSMA expression was restricted to smooth muscle cell linings of airways and larger-scale vasculature (**Figure 1c**). In contrast, bleomycin injured lungs revealed heterogeneously distributed regions of densified ECs and regions enriched for αSMA^+^ MFs both proximal to central bronchial airways as well as in the distal lung (**Figure 1d**).

**Figure 1:**
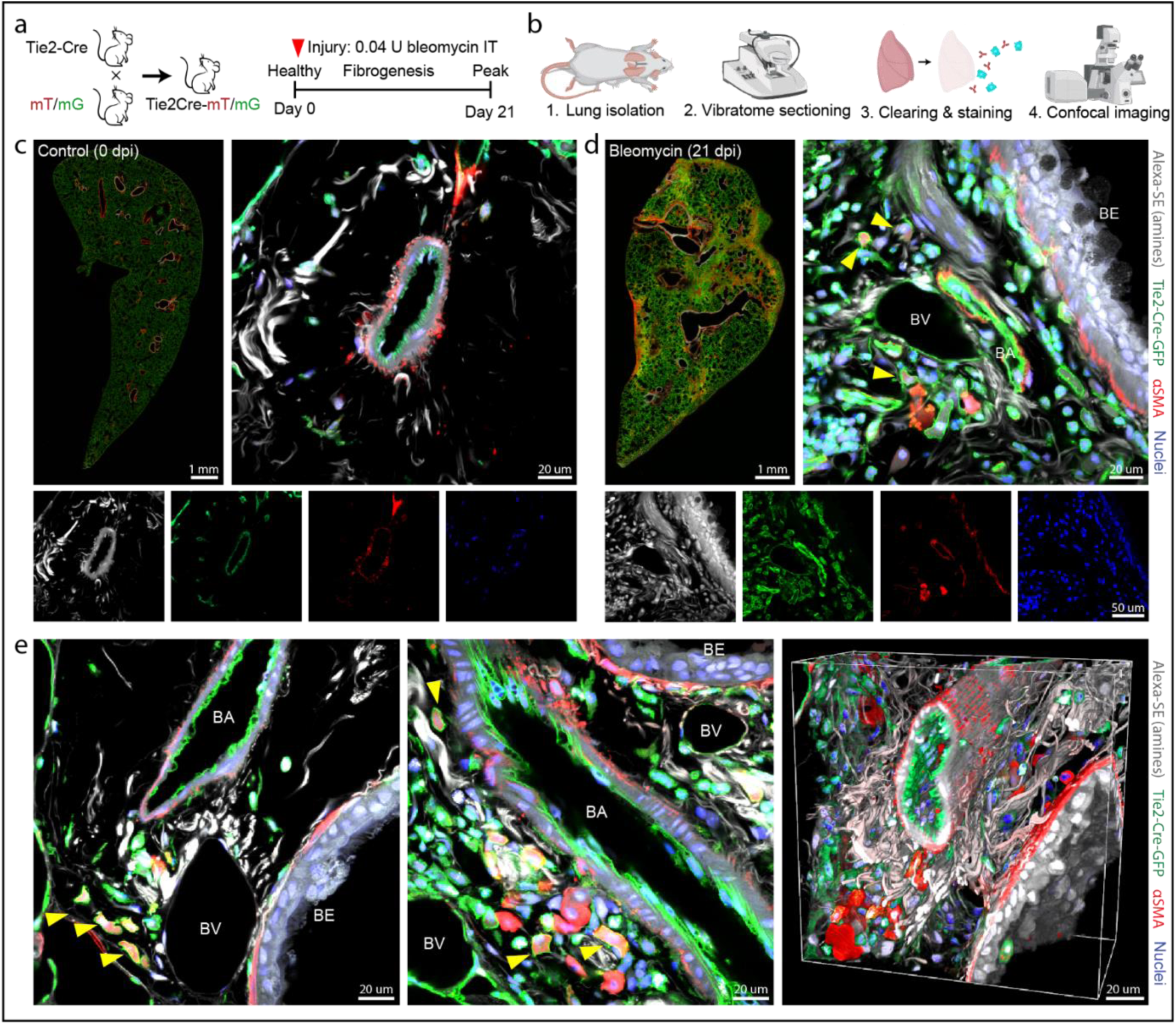
Emergence of aberrant ECs in the bleomycin lung injury model. a) Breeding strategy and study timeline for EC lineage tracing during bleomycin injury-induced lung fibrogenesis. b) Pipeline for 3D imaging of lung tissue. Representative low magnification whole-lobe tile-scan images and high magnification single z-slice images of peribronchial vasculature in (c) uninjured control lung and (d) bleomycin-treated lung tissues 21 days post-injury. e) Representative high magnification single z-slice images (left, middle) and 3D rendering (right) depicting cross-sectional and longitudinal views of bronchial vasculature. Yellow arrow heads mark individualized Tie2-Cre-GFP positive cells expressing αSMA. BA: bronchial arterioles, BV: bronchial venules, BE: bronchial epithelium.

Higher magnification confocal imaging revealed intriguing changes specific to the bronchial microvasculature (50-200 μm diameter, thereby excluding capillaries) and the perivascular ECM supporting these vessels (**Figure 1d,e)**. Bronchial arterioles (BA, identified by the presence of a α-SMA^+^ smooth muscle cell lining in large vessels proximal to bronchi) consistently possessed a surrounding fibrous matrix, as visualized by AFSE-labeling, which typically radiated outward from the BA. Bronchial veins (BV) appeared proximal to BAs, lacked smooth muscle cell investment, and typically possessed a markedly thinner vessel wall, possibly due to the potential for bronchial arteriolar blood flow return to the heart via the pulmonary venous system. Three weeks after bleomycin-induced injury, perivascular fiber density increased around BA/BVs and numerous GFP^+^ ECs were found to be embedded within this densified ECM. Interestingly, these ECs did not constitute multicellular sprouts typically associated with angiogenesis, but instead appeared as dispersed, individualized cells lacking intercellular connections. A subpopulation of these ECs found proximal to BAs/BVs were additionally α-SMA^+^ (Fig. 1d,e, yellow arrows), suggestive of EndMT, although high magnification imaging revealed a complete lack of α-SMA localization to stress fibers and instead diffuse staining throughout the cytosol. As macrophages have also been shown to express Tie2^39,40^, we immunostained for pan-monocyte/macrophage marker F4/80 and confirmed that the majority of GFP^+^ cells (including those that were dually GFP^+^ and αSMA^+^) were not macrophages **(Supplementary Figure 1)**.

Overall, these studies indicate that following lung injury, the density of perivascular matrix fibers increases surrounding arteriole/venule-scale bronchial vasculature and concurrently, a population of aberrant ECs that lack intercellular connections appear and adhere to the perivascular matrix. Whether these two observations are linked, and if so, how ECM fiber density may regulate EC signaling and behavior remain unknown. Given the high degree of spatiotemporal heterogeneity in tissue response to bleomycin-induced injury and the fact that there are no means to directly modulate ECM fiber density *in vivo,* we turned to *in vitro* biomaterial and microphysiological system (MPS) approaches to answer these questions.

### Heightened matrix fiber density induces formation of aberrant ECs in the absence of soluble cues

To test whether perivascular matrix fiber density modulates a switch between EC quiescence vs. activation and subsequent invasion, we integrated a previously established a multiplexed MPS containing perfusable arteriole/venule-scale microvessels^41–43^ with tunable, hybrid natural-synthetic hydrogel-fiber composites previously established by our group to model interstitial or stromal matrix^44–46^. Each MPS contains two parallel microchannels (Ø = 140 μm) fully embedded within a user-defined hydrogel, with each channel terminating in individually addressable media reservoirs **(Figure 2a)**. Here, we used fibrin (10 mg/ml), a naturally derived hydrogel commonly employed to model ECM during wound healing ^47^ which is known to leak from hyperpermeable vasculature and accrue in the extravascular space during lung injury and fibrosis^48^. One of each pair of channels was seeded with ECs, which self-assembled into a patent, arteriole/venule-scale parent microvessel overnight (16 hours post cell seeding) possessing VE-cadherin-enriched AJs and low vessel wall permeability **(Figure 2a and Supplementary Figure 2a-c)**^43^.

**Figure 2:**
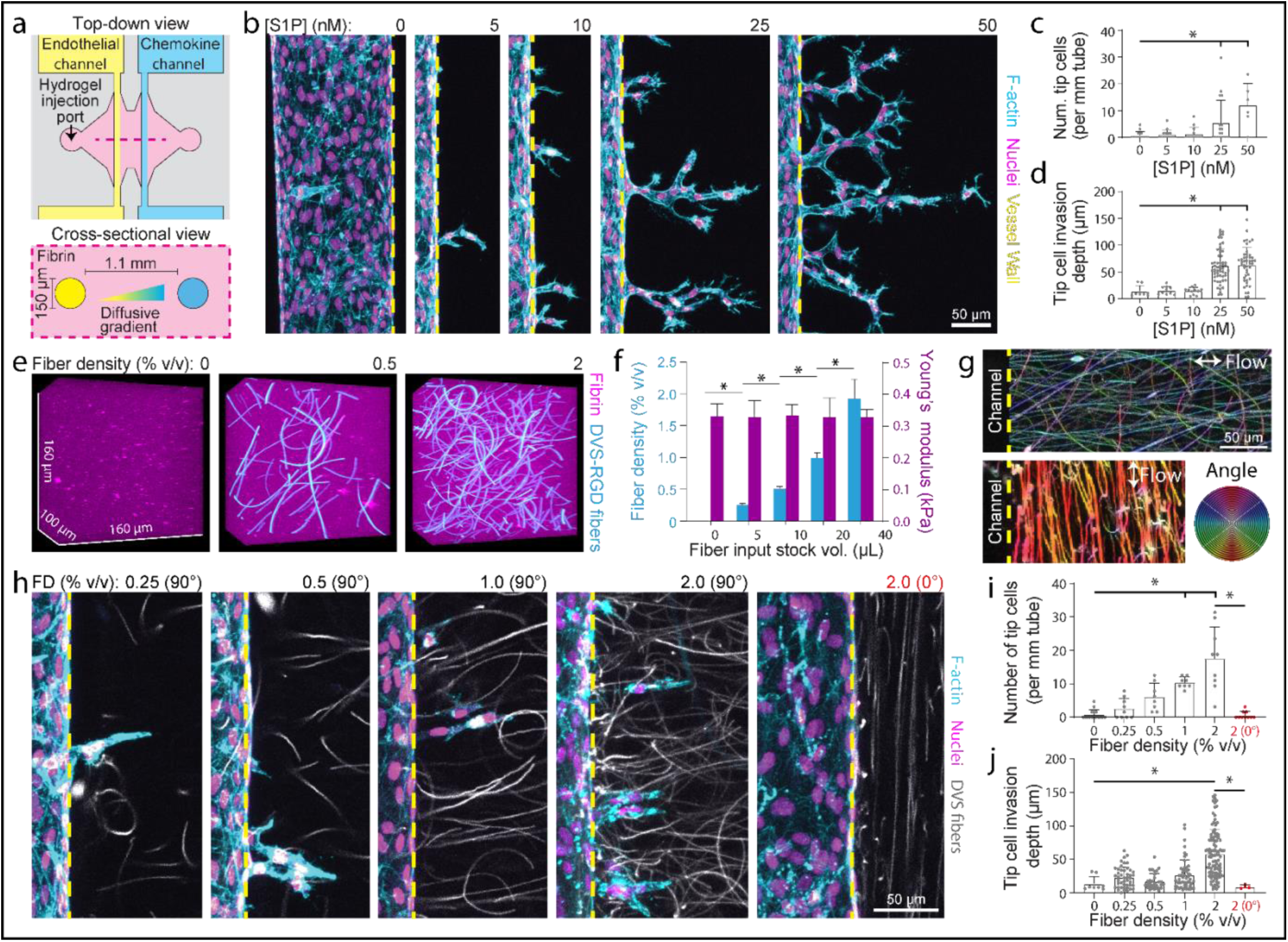
Heightened matrix fiber density induces spontaneous tip cell formation in the absence of soluble cues. a) Schematic of tip cell formation assay. b) S1P-induced tip cell formation over 4-days within 10 mg/ml fibrin hydrogels at indicated S1P concentrations. Nuclei (magenta), F-actin (cyan), yellow dashed lines indicate parent vessel edge. c-d) Quantification of number of tip cells and tip cell invasion depth as a function of [S1P] from (b). e) 3D rendering of fiber-embedded fibrin hydrogels. Synthetic DexVS fiber segments (cyan), fluorescently-labeled fibrin hydrogel (magenta). f) Quantification of fiber density and Young’s modulus as a function of input fiber stock volume. g) Flow-induced fiber alignment perpendicular (0°) or parallel (90°) to the long axis of the parent vessel. h) Fiber-induced tip cell formation (with no S1P added to the chemokine channel) over 4-days within 10 mg/ml fibrin hydrogels and indicated fiber density. Nuclei (magenta), F-actin (cyan), fibers (white), yellow dashed lines indicate parent vessel edge. i-j) Quantification of number of tip cells and tip cell invasion depth as a function of fiber density. Blue indicates 90° fiber alignment. All data presented as mean ± std.; * indicates a statistically significant comparison with P<0.05 (one-way analysis of variance).

Chemokines can be added to the adjacent, unseeded channel to generate a diffusive chemoattractant gradient that drives endothelial tip cell formation and ensuing angiogenic sprouting **(Figure 2a)**^41–43^. To confirm that this platform enables a robust, quantitative assessment of tip cell formation, we introduced a well-established EC chemoattractant, sphingosine-1-phosphate (S1P)^43,49^, to the chemokine channel at varying concentrations. Indeed, the number and invasion depth of invading tip cells increased as a function of S1P concentration and resulting gradient strength **(Figure 2b-d)**, establishing a robust assay and quantitative metrics for subsequent studies. Notably, our prior work demonstrates that the combination of a chemokine gradient (S1P) and mitogenic signals can activate tip cells which lead ensuing stalk cells to form perfusable capillaries, however changes in matrix density can engender single cell in lieu of multicellular sprout invasion^43,50^.

Individual microenvironmental cues presented by fibrin hydrogels are challenging to decouple and orthogonally tune (eg. hydrogel stiffness, adhesive ligand density, and porosity all vary as a function of fibrinogen density). Furthermore, our above characterization of increased density of perivascular fibers revealed matrix fibers with multi-micrometer diameters far greater than the <200 nM diameters typical of fibrin^51^ (**Figure 1**). Thus, we implemented our recently established composite hydrogel approach where a natural or synthetic bulk hydrogel can be combined with chemically and mechanically defined synthetic fiber segments that possess the geometry and mechanics of collagen fibrils prominent in the interstitial ECM of patients with idiopathic pulmonary fibrosis and other forms of interstitial lung disease^45^. Cell-adhesive fiber segments were generated via electrospinning of a synthetic polymer solution, dextran vinyl sulfone (DexVS), followed by segmenting fibers to defined lengths, and functionalization with the cell-adhesive peptide RGD. DexVS fiber segments were then incorporated into the fibrin hydrogel precursor solution at controlled v/v % to define the fiber density (0 – 2 v/v %) of the resulting hydrogel composite. Modulation of DexVS-RGD fiber density did not alter the stiffness of the bulk fibrin hydrogel as measured by AFM nanoindentation **(Figure 2e-f),** but likely influences local, cell-scale mechanics given that these fibers individually are considerably stiffer than bulk fibrin^42,52^. In prior work employing this composite approach, we demonstrated that heightened fiber density promotes myofibroblast activation and fibrogenic activity of encapsulated fibroblasts, in contrast to simply increasing the density or crosslinking/stiffness of an amorphous bulk hydrogel that lacks fibrous microstructure^45^. Here, we employed this approach to examine whether perivascular matrix fiber density modulates the activation of quiescent ECs into invasive tip cells.

Additionally, to model the matrix fiber alignment characteristic of the perivascular matrix observed in our *in vivo* studies (**Figure 1**), we adopted a flow-induced fiber alignment methodology previously utilized for aligning collagen fibrils within type 1 collagen matrices^53,54^. To generate fiber alignment perpendicular to the long axis of the parent vessel (90° alignment) as we observed *in vivo*, the hydrogel precursor solution was injected orthogonal to inserted channel-molding needles to generate a fluid flow profile that aligns fibers radially with respect to the eventual vessel **(Figure 2g)**. To align fibers parallel to the long axis of the parent vessel (0° alignment), hydrogel precursor solution was first injected into the device and subsequent insertion of acupuncture needles resulted in fluid flow profiles that aligned the fibers in the direction of needle insertion (ie. parallel to eventual parent vessels). Prior to cell seeding, the microchannel was coated with basement membrane proteins (Matrigel) to establish equivalent initial matrix ligand composition and topography for ECs adhering to microchannel walls, despite variations in fiber alignment and density in the surrounding matrix. Indeed, 16 hours after EC seeding, parent vessels in non-fibrous control vs. fibrous fibrin hydrogels did not differ in vessel diameter, cell density, or permeability as assessed by fluorescently labeled dextran diffusion across the vessel wall **(Figure 4a,b)**.

To our surprise, over extended culture durations but in the absence of an exogenous angiogenic or chemotactic gradient, microvessels with as a function of perivascular fiber density (FD) demonstrated increases in EC activation and invasion potential, as quantified by the number of ECs divesting from parent vessels and their invasion distance from the parent vessel wall after 3 days of culture **(Figure 2h-j)**. Many of these ECs invaded as individualized cells, and so we termed them aberrant tip ECs (ATECs) given their comparable morphology to tip ECs typical of angiogenic sprouts but evident lack of intercellular adhesion to ensuing stalk ECs. The formation of ATECs proved highly dependent upon fiber orientation with respect to the parent vessel, as an equivalently high density (2% v/v) of fibers, but aligned parallel to the parent vessel axis (0° alignment), resulted in virtually no EC invasion **(Figure 2h-j)**. These results strikingly demonstrate that both fiber density and alignment of perivascular matrix fibers dictate the activation of ECs and their subsequent invasion into the surrounding matrix, drawing clear parallels to prior work on EMT and epithelial cell migration in the context of cancer metastasis^55,56^.

The studies above employed human umbilical vein ECs as a model EC due to their frequent use throughout the field of endothelial cell biology. To assess whether ECs specific to tissues known to be susceptible to fibrosis would similarly respond to densified, radially aligned fibers surrounding an arteriole/venule –scale vessel, we performed identical studies with primary human liver, lung, and dermal microvascular ECs. Heightened fiber density led to increased ATEC formation and invasion in liver and lung microvascular ECs **(Supplementary Figure 3a-c)**. However, dermal microvascular ECs did not activate or invade to any degree. While our synthetic matrix-like fibers are chemically and mechanically defined, and thus highly tunable, their dextran-based composition is non-native to mammalian tissues; thus, we sought to confirm whether a similar phenomenon occurred in response to natural collagen fibers, given their ubiquitous presence in fibrotic tissue. To do so, we adopted a recently established approach to generating larger collagen fibers that possess comparable diameters to collagen fibers observed *in vivo* (Ø > 1 μm), in contrast to those comprising traditionally prepared collagen hydrogels (Ø = 250-500 nm)^45,54^. Collagen fibers were isolated and suspended within fibrin hydrogels and identical studies as described above demonstrated that collagen fiber density also induced ATEC formation **(Supplementary Figure 4)**. Finally, as greater than capillary-scale microvasculature normally possesses supporting mural cells, we additionally performed studies incorporating human mesenchymal stem cells (MSC) to generate mural cell-invested parent vessels ^57^. Despite the presence of mural cells, heightened fiber density induced ATEC formation and interestingly led to mural cell divestment and polarized migration into the surrounding matrix as compared to non-fibrous controls **(Supplementary Figure 5)**.

### ATEC formation due to increased interstitial fiber density involves matrix mechanosensing and EndMT

EndMT has been previously implicated in the transition of quiescent vessel-lining ECs into invasive tip cells^11,16,58^ and in separate work, associated with EC mechanosensing of stiffened ECM^6^. However, prior studies primarily relied on EC cultures on 2D substrates with tunable stiffness. In contrast, angiogenesis is inherently a 3D process involving the proteolytic invasion of ECs through a surrounding fibrous ECM. Thus, we examined whether ECM fiber density similarly influenced EC mechanosensing and EndMT in our 3D model. Invading ATECs possessed thinner AJs as evidenced by VE-cadherin immunostaining, in contrast to the more robust maintenance of AJs in adjacent ECs composing the parent vessel (**Figure 3a**). Supporting an EndMT-mediated activation of tip cells, SNAI1 (Snail) immunostaining revealed heightened cytosolic and nuclear SNAI1 localization in invading ECs. Consistent with these findings, ECs cultured on fibrous substrates increased their expression of EndMT-associated genes (*SNAI1, SNAI2, CDH2*), while decreasing VE-cadherin expression in contrast to controls on biochemically identical surfaces lacking fibrous topography (**Supplemental Figure 6**). YAP immunostaining revealed similar expression levels and distribution, with ATECs possessing higher overall levels of YAP and comparable distributions between cytosolic and nuclear compartments, in contrast to parent vessel ECs where YAP was clearly excluded from the nucleus (**Figure 3b-c**).

**Figure 3:**
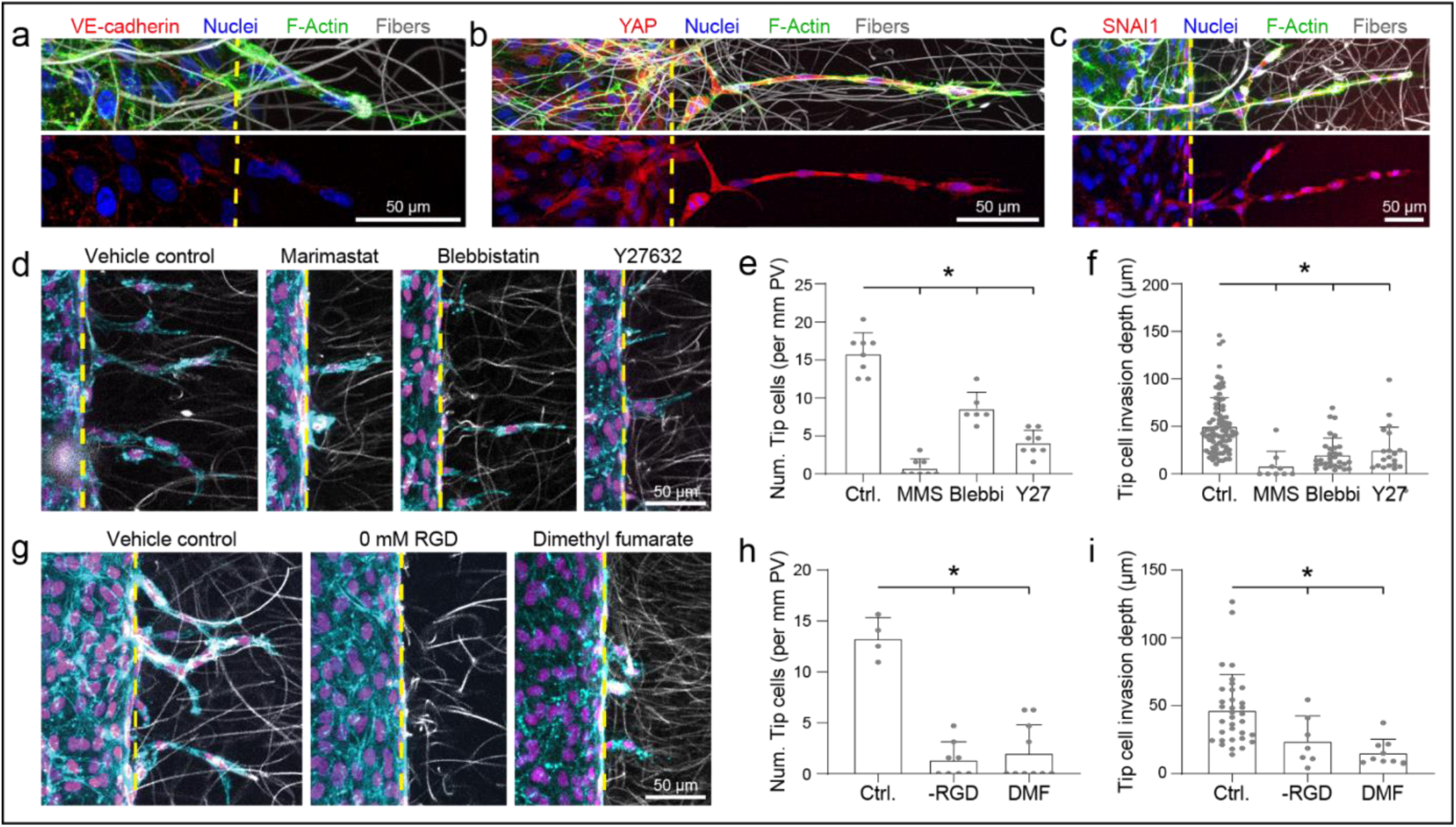
Fiber-mediated ATEC activation and invasion share signaling and cellular functions previously implicated in EndMT and angiogenic tip cells. a-c) Representative images depicting the differential localization of immunostained VE-cadherin (a), YAP (b), and SNAI1 (c) invading ECs vs. parent vessel ECs within after 4 days of culture. Yellow dashed lines indicate parent vessel edge. d-f) Representative images (d) of vehicle control, marimastat, blebbistatin, or Y27 treated microvessels within FD 2% hydrogels after 4 days of culture, along with corresponding quantification of the number of invading ECs (f) and their invasion depth (g). g-i) Representative images (g) of vehicle control microvessels with RGD-functionalized fibers (FD 2%), control microvessels with fibers lacking RGD functionalization (FD 2%), and dimethyl fumarate treated microvessels with RGD-functionalized fibers (FD 2%) after 4 days of culture and corresponding quantification of the number of invading ECs (h) and their invasion depth (i). All data presented as mean ± std.; * indicates a statistically significant comparison with p < 0.05 (two-sided student’s t-test compared to control condition).

**Figure 4:**
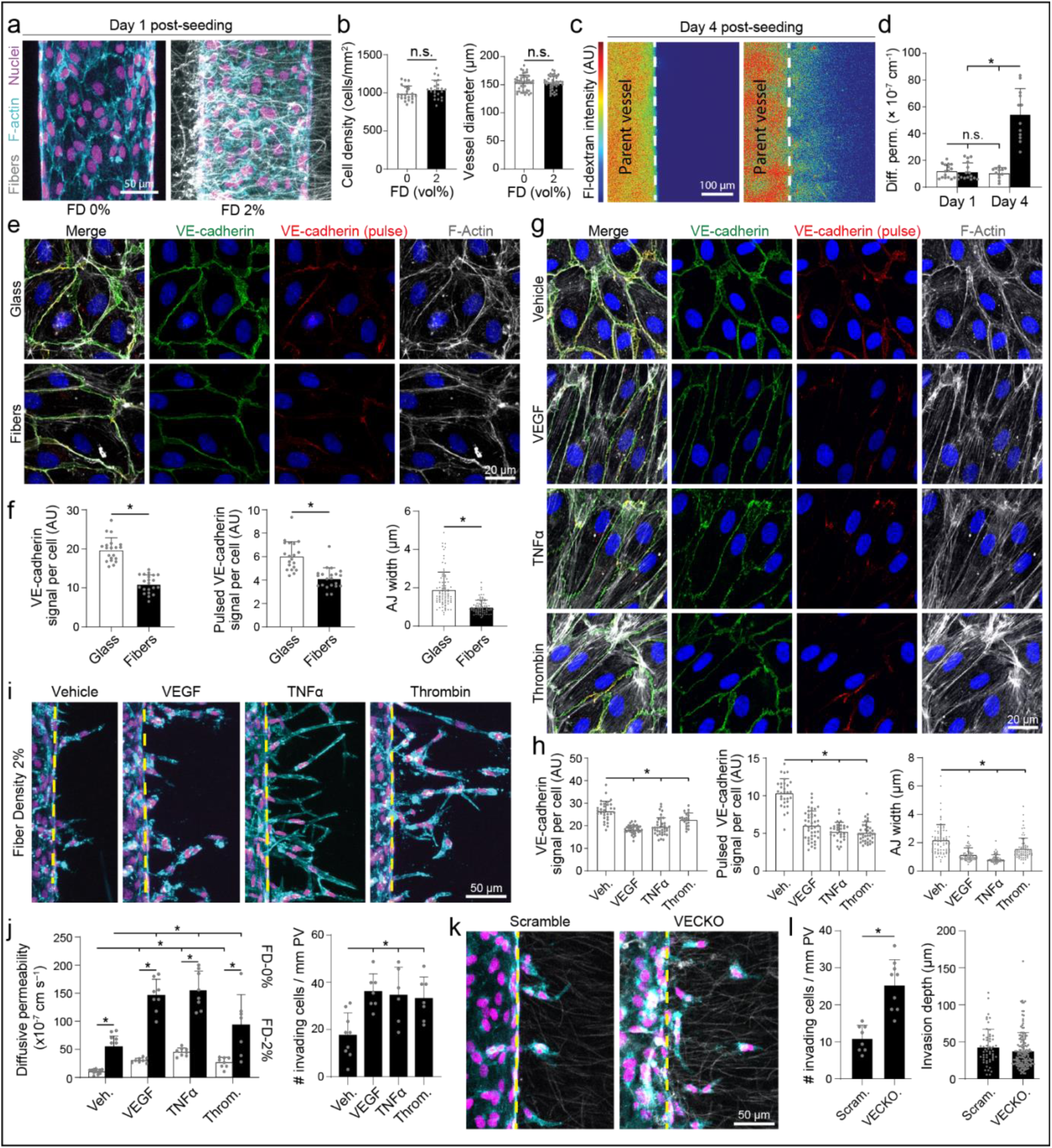
VE-cadherin destabilization increases vessel permeability and ATEC activation. a-c) Representative maximum intensity projection images of microvessels with or without FD 2% cell-adhesive fibers after one day of culture along with corresponding quantification (b) of vessel wall cell density and microvessel diameter. c) Representative single z-slice (at mid-plane of microvessel) following dextran perfusion on day 4 of culture. d) Quantification of vessel wall diffusive permeability of microvessels cultured for 1 or 4 days in FD 0% or 2% hydrogels. e-f) Representative images (e) of endothelial cell monolayers cultured for 2 days on glass coverslips or electrospun DexVS fiber matrices along with corresponding quantifications (f) of VE-cadherin signal per cell, pulsed VE-cadherin signal per cell, and adherens junction (AJ) width as determined by segmentation of VE-cadherin^+^ cell-cell junctions. g-h) Representative images (g) of endothelial cell monolayers cultured for 2 days on fibrous matrices treated with vehicle, VEGF (50 ng/ml), TNFα (50 ng/ml), or thrombin (2 U/ml) along with corresponding quantifications (h) of VE-cadherin signal per cell, pulsed VE-cadherin signal per cell, and adherens junction (AJ) width as determined by segmentation of VE-cadherin^+^ cell-cell junctions. i-j) Representative maximum intensity projection images (i) of microvessels with FD 2% cell-adhesive fibers treated with vehicle, VEGF (50 ng/ml), TNFα (50 ng/ml), or thrombin (2 U/ml) after 4 days of culture along with quantification (j) of vessel wall diffusive permeability and number of invading ECs. Yellow dashed lines indicate parent vessel edge. k-l) Representative maximum intensity projection images (k) of microvessels with FD 2% cell-adhesive fibers composed of scramble control of VE-cadherin KO ECs after 4 days of culture along with corresponding quantification (l) of the number of invading ECs and their invasion depth. All data presented as mean ± std.; * indicates a statistically significant comparison with P<0.05 (b, d-f, i: two-sided student’s t-test, j-l, n, p: one-way analysis of variance).

Towards identifying critical cellular requirements for the activation and invasion of ATECs, we utilized a panel of pharmacologic inhibitors targeting mechanosensing and EndMT-associated cell functions. We tested requirements for matrix proteolysis and actomyosin contractility, both of which have been previously shown to be gained by ECs that have undergone EndMT^10^. Treatment with marimastat, a broad-spectrum inhibitor of MMP-mediated proteolysis, decreased both the number of ATECs and their invasion depth, as ECs were unable to degrade the surrounding matrix and generate sufficient space for invasion **(Figure 3d-f)**. Treatment with a myosin II inhibitor (blebbistatin) or ROCK inhibitor (Y-27632) resulted in decreased ATEC formation, in line with the premise that actomyosin contractility is a key requirement for cell migration in 3D and matrix mechanosensing **(Figure 3d-f)**. As the protrusions of invading ECs were observed to extend along fibers (**Figure 3a-c**), we next tested whether direct integrin-mediated adhesion to fibers was critical for ATEC invasion. Indeed, identical studies performed with fibers lacking RGD functionalization resulted in reduced ATEC numbers and invasion distances **(Figure 3g-i)**, indicating that direct EC engagement to matrix fibers is a requirement for ATEC formation. Lastly, recent studies in addition to the studies described above have demonstrated that fibrous matrices can modulate YAP/TAZ signaling in EC monolayers to promote cell migration^59^. Providing further support for the involvement of YAP/TAZ signaling in EC activation and invasion in this setting, treatment with dimethyl fumarate, a potent although non-specific inhibitor of the YAP/TAZ signaling pathway^60^, resulted in decreased ATEC formation **(Figure 3g-i)**. Overall, these studies establish that proteolysis, actomyosin contractility, cell-matrix adhesion, and mechanosensing through YAP/TAZ are all required for the transition from EC quiescence to activation and invasion into a surrounding fibrous matrix.

### Fiber-induced AJ destabilization decreases vessel barrier function and promotes ATEC invasion

Taken together, our results demonstrate that ECs within arteriole/venue-scale vessels sense heightened perivascular fiber density, which engenders EndMT signaling to contribute to ATEC formation from an otherwise quiescent endothelium. Although barrier function was equivalent on day 1, three additional days of culture led to a 5-fold increase in permeability for vessels exposed to a heightened perivascular fiber density as compared to controls, which retained low permeability throughout the same duration of study **(Figure 4a-d)**. As increased vessel permeability and EndMT signaling have both associated with the destabilization of VE-cadherin-based AJs^61,62^ we next examined whether EC adhesion to a fibrous topography directly modulated AJs.

We assessed AJs via VE-cadherin immunostaining in confluent EC monolayers cultured on ‘flat’ substrates (lacking fibrous topography) as compared to ∼2D substrates composed of identical fibers deposited on glass coverslips ^52,63^. In line with our observations of decreased vessel barrier function as determined by permeability measurements, EC monolayers on fibrous substrates were characterized by decreased overall VE-cadherin intensity at AJs and VE-cadherin-containing AJ width **(Figure 4e, f)**. We next examined relative VE-cadherin stability as a function of fibrous topography by live pulse-labeling with a non-perturbing, fluorescently tagged anti-VE-cadherin antibody that binds to the extracellular domain of VE-cadherin. Relatively higher intensity of pulse-labeled VE-cadherin would be indicative of slower VE-cadherin turnover^64^. EC monolayers on a fibrous topography possessed lower pulsed anti-VE-cadherin antibody intensity at AJs as compared to non-fibrous ‘flat’ controls **(Figure 4e, f)**, suggesting that decreased junctional VE-cadherin observed in EC monolayers on fibrous topography may be due in part to increased turnover rates at which VE-cadherin is internalized from AJs.

VE-cadherin abundance at AJs appeared to be modulated by EC interactions with a fibrous topography, and previous studies have implicated AJ stability vs. disassembly to be critical determinants in EC quiescence vs. activation. To test whether destabilization of AJs promotes fiber-induced ATEC formation, we next examined ATEC formation after treatment of parent vessels with three well-established vessel permeability agonists – VEGF, TNFα, and thrombin ^65–67^. We first established concentrations of VEGF, TNFα, or thrombin that led to decreased VE-cadherin AJ localization, AJ width, and pulsed VE-cadherin signal intensity when delivered to EC monolayers cultured on ‘flat’ substrates **(Figure 4g,h)**. As expected, treatment of EC microvessels in both matrix conditions with optimized concentrations of VEGF, TNFα, or thrombin resulted in increases in vessel wall permeability. However, we observed that heightened perivascular fiber density in combination with any of the three permeability agonists resulted in increased ATEC formation **(Figure 4i,j).** To directly test the role of VE-cadherin in gating fiber-induced ATEC formation, we examined an extreme scenario of ECs completely lacking VE-cadherin by complete knockout of VE-cadherin (VE-cadherin knockout, VECKO) via CRISPR-Cas9 editing. Microvessels formed from VECKO ECs in matrices with heightened fiber density revealed heightened EC invasion as compared to scrambled gRNA controls **(Figure 4k,l)**. These results collectively demonstrate that AJ destabilization resulting from EC adhesion to matrix fibers results in destabilized VE-cadherin adherens junctions which potentiates ATEC formation and invasion.

### Transcriptomic analysis of ATECs reveals EndMT signaling and a proinflammatory phenotype

We next explored the EC transcriptome as a function of EC organization (ie. multicellular microvessels vs. isolated cells) and engagement with a 3D fibrous topography with the goal of characterizing differential gene expression and biological processes characteristic of ATECs. For multicellular samples, mRNA was harvested from pooled MPS microvessels cultured for 3 days, alongside a control consisting of a confluent monolayer on fibrin (**Figure 5a-c**). Given the limited number of ATECs formed per MPS microvessel (∼50) and our inability to isolate them from ECs remaining part of the microvessel, we modeled ATECs (ie. isolated ECs adhering to matrix fibers) at sufficient quantities by encapsulating individualized cells in fibrous (FD 2% v/v) fibrin hydrogels and included a non-fibrous (FD 0% v/v) condition with the goal of segregating effects of 3D encapsulation from those arising from cell adhesion to matrix fibers. Encapsulated ECs cultured for 3 days (prior to harvest for transcriptomic analysis) in matrices with heightened fiber density possessed greater projected spread areas and longer protrusions compared to non-fibrous controls (**Figure 5a**), in line with our prior studies demonstrating that cell-adhesive fibers promote EC spreading^68^. Although ECs in this condition lacked a shared orientation due to the random orientation of fibers in bulk-casted gels, individually these cells adopted a uniaxial, spindle morphology and extended protrusions along fibers, identical to invading ATECs in earlier MPS studies **(Figure 2)**.

**Figure 5:**
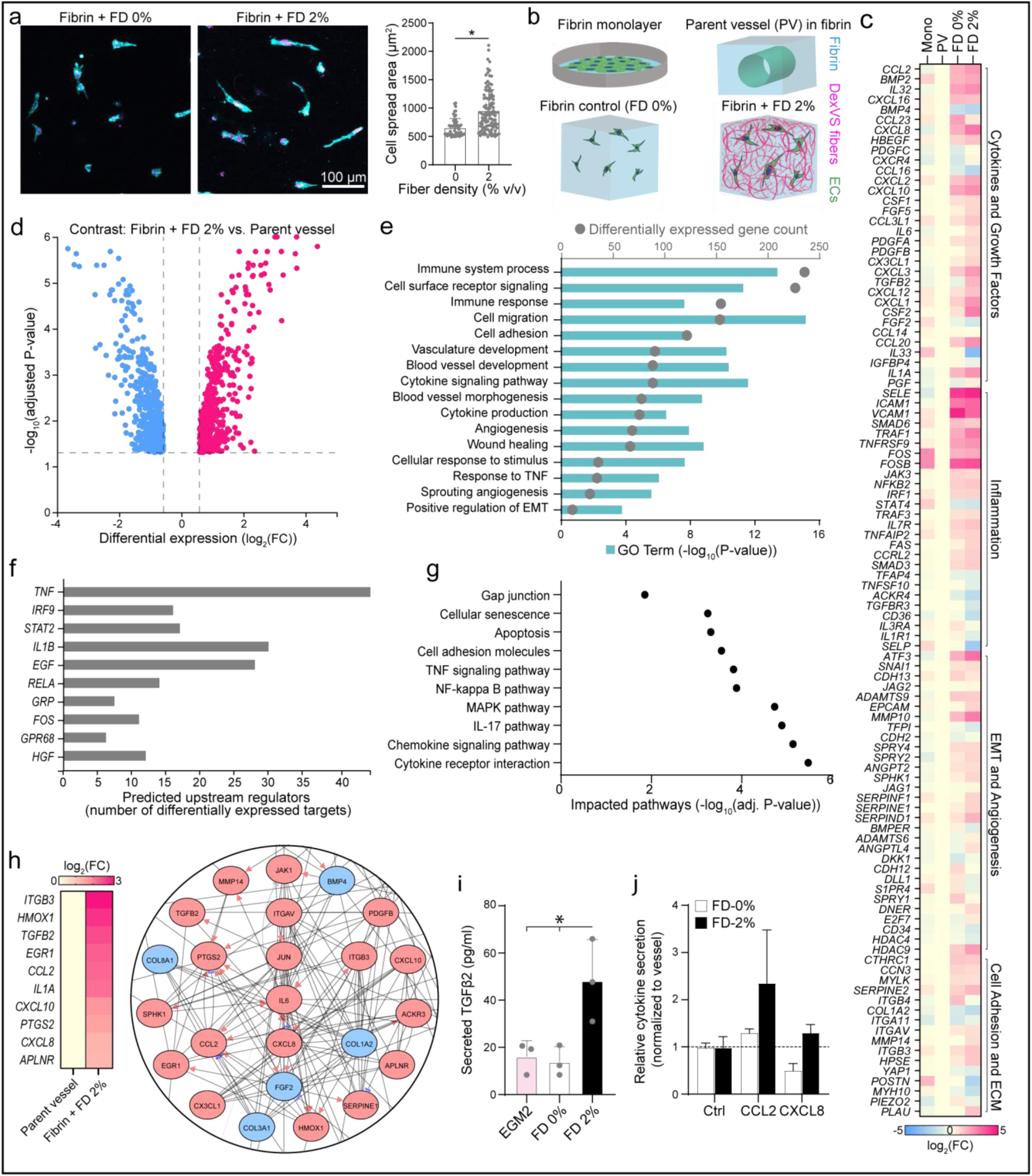
Transcriptomic analysis of ATEC suggests an inflammatory and pro-fibrotic secretory phenotype. a) Representative maximum intensity projection images of ECs embedded within control (FD 0%) or fibrous (FD 2%) hydrogels cultured over 3 days to model ATECs at sufficient numbers for transcriptomic analysis and corresponding quantification of EC spread area. b) Schematic depiction of conditions for transcriptomic analysis. c) Expression of several curated gene categories relative to parent microvessels after 3 days of culture. d) Volcano plot visualizing differentially expressed genes in ATECs as compared to parent microvessel ECs after 3 days of culture. e) P-value and number of differentially regulated genes for Gene Ontology terms comparing ATECs in FD 2% to parent microvessel ECs. h) Curated genes of interest differentially expressed in FD 2% vs monolayer conditions (left). Predicted interactome for differentially expressed genes relevant to fibrosis (right). i) Microfluidic ELISA quantification of TGF-β2 secretion. j) Cytokine antibody membrane array detection of CCL2 and CXCL8. b, i: Data presented as mean ± std; * indicates statistically significant comparison with P<0.05 (b: two-sided student’s t-test. i: one-way analysis of variance).

Comparing confluent 2D EC monolayers adhering to fibrin to ECs isolated from MPS microvessels, relatively few differentially regulated genes were identified **(Figure 5c),** indicating comparable gene expression programs despite fluidic perfusion in MPS cultures and minor differences in substrate curvature. We next focused our analysis on whether single ECs engaging a matrix with heightened fiber density adopted a transcriptomic signature that might promote fibrosis-associated signaling as compared to quiescent ECs within multicellular microvessels.

In contrast, ECs cultured in 3D in FD 0% and 2% fibrin hydrogels possessed 294 and 1071 differentially expressed genes, respectively **(Figure 5d)**. Heightened fiber density (FD 2% v/v), which gave rise to ATECs in earlier MPS experiments (**Figures 2-4**), significantly changed Gene Ontology terms associated with cell migration, cell adhesion, angiogenesis, and positive regulation of EMT, all of which involved genes implicated in tip cell activation, migration, or function (**Figure 5e)**. Supporting our earlier observations of increased SNAI1 immunostaining in ATECs (**Figure 3b**) and gene expression resulting from EC adhesion to a fibrous topography (**Supplementary Figure 6**), *SNAI1* gene expression increased along with several other EndMT-associated genes such as *MMP14* and multiple TGF-β-associated genes (*TGFB2, BMP2, SMAD3*) **(Figure 5c and Supplementary Figure 7)**. Expression of *CTHRC1*, recently established as a unique marker of disease-associated myofibroblasts in the heart and lung, was comparably increased in both FD 0% and FD2%, although we noted decreased expression of genes associated with matrix secretion and remodeling (*COL1A2* and *POSTN)*, two well-accepted markers of myofibroblasts, suggesting that although ECs may undergo EndMT, transition to *bona fide* myofibroblast may require additional soluble and/or physical factors not present in these studies ^7,10,69^. Consistent with these findings, only a sub-population of lineage-traced ECs were also αSMA^+^, and αSMA within these cells did not localize to stress fibers but instead appeared uniformly diffuse throughout the cytosol (**Figure 1**).

Interestingly, we observed TGF-β2 gene expression to be uniquely upregulated in ECs cultured in matrices with heightened fiber density **(Figure 5)**. As the family of TGF-β growth factors broadly are potent mediators of fibrosis via influencing fibroblast-myofibroblast transition and EndMT, we employed a microfluidic ELISA assay to quantitatively measure TGF-β2 secretion levels as a function of EC engagement to matrix fibers. TGF-β2 secretion from ECs encapsulated in matrices with heightened fiber density was 3-4 fold higher than nonfibrous controls, which were comparable to baseline levels of TGF-β2 within the serum component of EGM2 **(Figure 5i)**. Furthermore, we utilized a cytokine antibody membrane array to detect whether other secreted cytokines were elevated in response to heightened fiber density. An inflammatory cytokine immunoblot array indicated elevated levels of MCP1 (CCL2) and IL8 (CXCL8) secreted by ATECs compared to parent vessel ECs **(Figure 5j and Supplementary Figure 8a-b)**. In line with these findings, both CCL2 and CXCL8 were significantly upregulated at the transcriptomic level **(Figure 5h)**. Taken together, our data demonstrates ATECs express genes associated with EndMT markers and angiogenesis, as well as pro-inflammatory and pro-fibrotic cytokines suggestive of a disease propagating phenotype that is distinct from *bona fide* myofibroblasts.

### A VE-cadherin–TGF-βR2 interaction gates fiber-induced TGF-β signaling in ECs

Our data suggest that EC adhesion to a fibrous topography destabilizes AJs thereby enabling EndMT signaling and ATEC formation (**Figure 4**) and that ATECs secrete TGF-β2 (**Figure 5**). As AJ and VE-cadherin turnover have previously been shown to regulate EC behavior in response to growth factors such as VEGF^70^, we next sought to identify potential molecular mechanisms by which VE-cadherin stability at AJs might regulate TGF-β signaling in ECs. To do so, we employed proximity ligation mass spectrometry (BioID) to profile proteins that associate with VE-cadherin in quiescent, confluent monolayers reflective of the adhesive state of ECs within quiescent microvessels (**Figure 6a,b**), supported by the above finding that EC monolayers and MPS microvessels are transcriptionally quite similar. In quiescent 2D EC monolayers, analysis of the most abundant proteins identified via mass spectrometry expectedly revealed prominent associations with core AJ plaque proteins including α, β, γ, and δ-catenins. However, among the most abundant (top 15) VE-cadherin interactors, we unexpectedly identified a prominent association between VE-cadherin and the receptor tyrosine kinase transforming growth factor β receptor 2 (TGF-βR2) (**Supplementary Figure 9**). As our transcriptomic and functional analyses associate EndMT signaling with ATECs, and TGF-β signaling has been established to drive EndMT, we next investigated whether this putative VE-cadherin-TGF-βR2 interaction is modulated by EC engagement to a fibrous topography.

**Figure 6:**
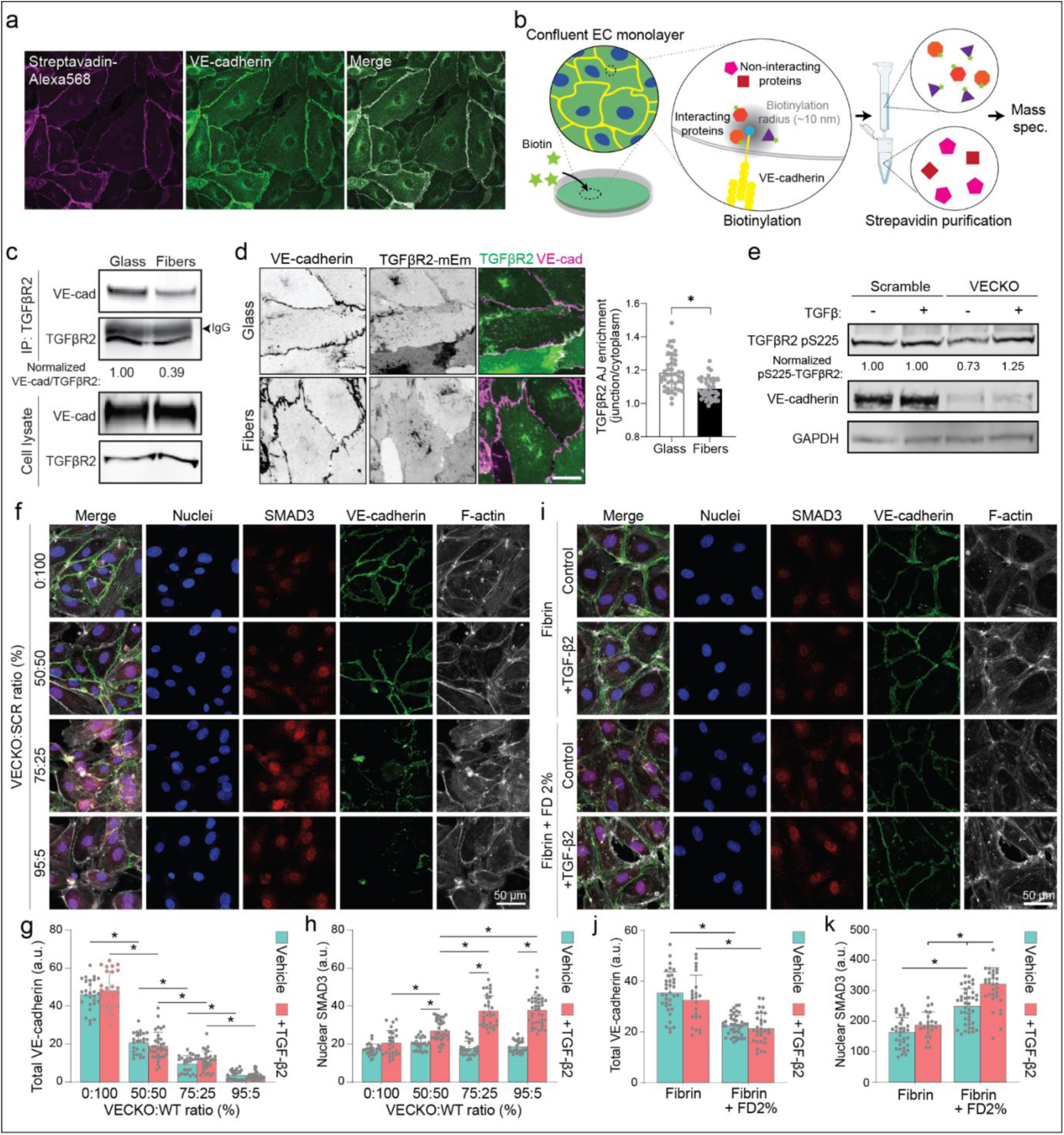
EC engagement to fibrous topography modulates TGF-β signaling via VE-cadherin–TGF-βR2 dependent interactions. a) ECs stably expressing VE-cadherin-BioID incubated were incubated biotin in media and then stained for VE-cadherin and biotinylated proteins (Streptavidin-AF568). b) Schematic of VE-cadherin-BioID mass spectrometry screen to identify interactors of VE-cadherin c) Western blot micrograph of VE-cadherin co-immunoprecipitation stained for TGF-βR2 from ECs cultured on flat glass or fibrous substrates. Blot representative of one independent experiment. d) Representative images of ECs expressing TGF-βR2-mEmerald cultured on flat substrates or fibrous topographies immunostained for VE-cadherin along with quantification TGF-βR2 localization to VE-cadherin AJs. e) Western blot of scrambled control or VECKO EC lysates treated with or without TGF-β2 ligand (10 ng/mL). Blot representative of two independent experiments. f-h) Representative images (f) and quantifications of total VE-cadherin intensity (g) and nuclear SMAD3 (h) as a function of varied ratios of scrambled control ECs and VECKOs. i-k) Representative images (i) and quantifications of total VE-cadherin intensity (j) and nuclear SMAD3 (k) as a function of EC adhesion to pure fibrin (FD 0%) or fibrin + FD 2%.

To test whether TGF-βR2 and VE-cadherin association is influenced by EC engagement to matrix fibers, EC monolayers were cultured on ‘flat’ glass substrates or substrates with fibrous topography, lysed, and VE-cadherin interactors were analyzed by co-immunoprecipitation. We confirmed that VE-cadherin co-immunoprecipitates with TGF-βR2 and that less TGF-βR2 associates with VE-cadherin in ECs cultured on fibers compared to those cultured on flat substrates (**Figure 6c**), suggesting that EC engagement to a fibrous topography attenuates this VE-cadherin–TGF-βR2 interaction. Corroborating this finding, expression of an mEmerald-fusion tagged TGF-βR2 in ECs on flat vs. fibrous topographies showed decreased enrichment of TGF-βR2 at AJs in ECs engaging a fibrous topography (**Figure 6d**). To test whether VE-cadherin interactions with TGF-βR2 modulated the EC response to TGF-β ligands, we next assayed TGF-βR2 phosphorylation^71^ in scrambled control vs. VECKO EC monolayers. Stimulation with TGF-β ligand did not alter TGFβ-R2 phosphorylation in control EC monolayers. In VECKO EC monolayers, basal TGF-βR2 phosphorylation is lower than control ECs; however, in contrast, TGF-βR2 phosphorylation markedly increases in response to stimulation with TGF-β (**Figure 6e**). Taken together, we identify a VE-cadherin–TGF-βR2 interaction that is disrupted by heightened fiber density, in turn regulating the phosphorylation state of TGF-βR2 in the presence of TGF-β ligands.

To test whether TGF-β signaling is modulated by VE-cadherin and AJ stability, we next performed mosaic studies with scramble control and VECKO ECs at varying ratios to modulate VE-cadherin localization to AJs independent of cell density. To assess TGF-β signaling, we treated cells with TGF-β2, as our prior findings demonstrated that ATECs secrete this TGF-β ligand (**Figure 5i**), and quantified nuclear SMAD3 localization, as TGF-β2 binding to TGF-β receptors leads to downstream nuclear translocation of SMAD2/3/4 transcription factor complexes ^72^. Indeed, increased ratios of VECKO:control ECs resulted in decreased VE-cadherin intensity at AJs and corresponding increases in SMAD3 nuclear localization **(Figure 6f-h)**. Additionally, we employed a scratch wound assay to vary AJs within the same samples where ECs at the scratch margin should possess less ECs than those fully surrounded by neighboring cells. At the scratch wound margin, ECs decreased VE-cadherin and increased nuclear SMAD3 compared to regions distanced from the scratch which were more cell dense with increased VE-cadherin and decreased nuclear SMAD3 **(Supplementary Figure 10a-c)**. Lastly, we performed studies comparing control, non-fibrous matrices with fibrous (FD 2%) matrices and found that fibrous topography without addition of TGF-β2 led to increased nuclear pSMAD3 which was further increased with TGF-β2 treatment **(Figure 6i-k)**. Altogether, these studies demonstrate that VE-cadherin inhibits downstream TGF-β-signaling via an interaction with TGFβ-R2, resulting in decreased nuclear SMAD3 expression, and this interaction is disrupted upon VE-cadherin destabilization by EC adhesion to fibrous ECM.

### TGFβ2-induced apoptosis underlies microvasculature rarefaction in late-stage fibrosis

Our transcriptomic and secretomic analyses implicate ATECs as a potential fibrosis-propagating EC phenotype via inflammatory cytokine and TGF-β2 secretion, which may drive fibrosis through action on other tissue resident cells (i.e. macrophages, fibroblasts, endothelium). To understand how increased ATEC-secreted TGF-β2 may influence arteriole-scale vasculature, we next treated parent vessels in control and fibrous hydrogels with exogenous TGF-β2. TGF-β2 treatment significantly reduced parent vessel cell density in both control and fibrous conditions through TGF-β2-induced apoptosis^73^, as compared to untreated vessels. However, TGF-β2-induced apoptosis was enhanced in parent vessels in fibrous conditions, as evidenced by ECs expressing cleaved-caspase-3 **(Figure 7a)**. In line with these findings, TUNEL staining confirmed evident EC apoptosis in bronchial arterioles and venules of bleomycin-dosed lungs 3 weeks after injury, in stark contrast to saline-treated controls **(Figure 7b)**. Cells resident to airways or alveolar regions demonstrated no indication of apoptosis, although abundant dual positive GFP/αSMA ECs were present.

**Figure 7:**
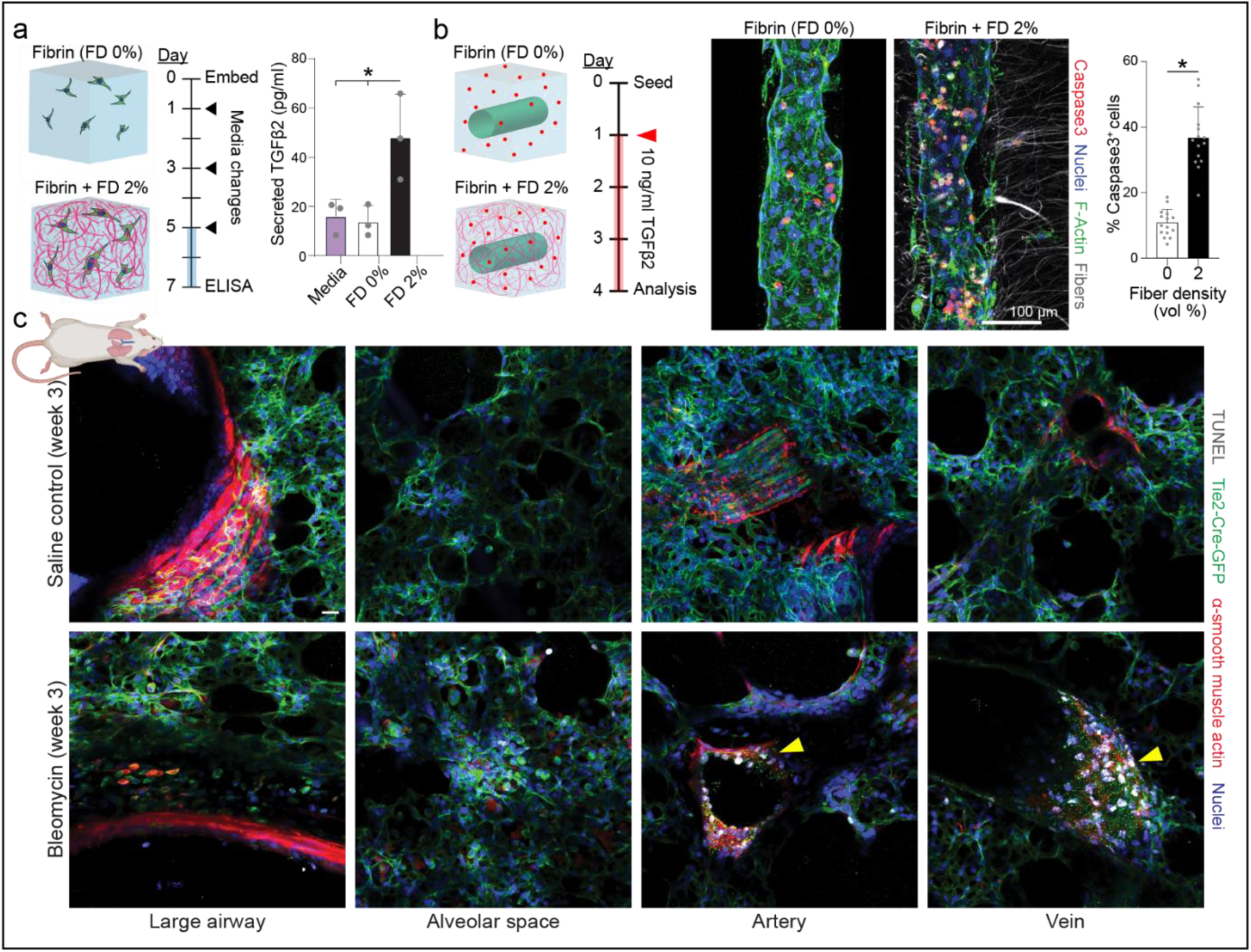
EC engagement to a matrix with heightened fiber density potentiates apoptosis *in vitro* and *in vivo.* a) Schematic of study conditions and timeline for assessing TGF-β2 secretion from ECs encapsulated in fibrin with or without heightened fiber density. b) Quantification of TGF-β2 secretion in culture media was performed on day 7 via microfluidic ELISA. c) Schematic of study conditions and timeline to assess the impact of heightened perivascular fiber density on TGF-β2-mediated apoptosis. d) Representative images of large airways, alveolar space, arterioles, and venules from saline control and bleomycin treated mouse lungs at 3 weeks.

**Figure 8:**
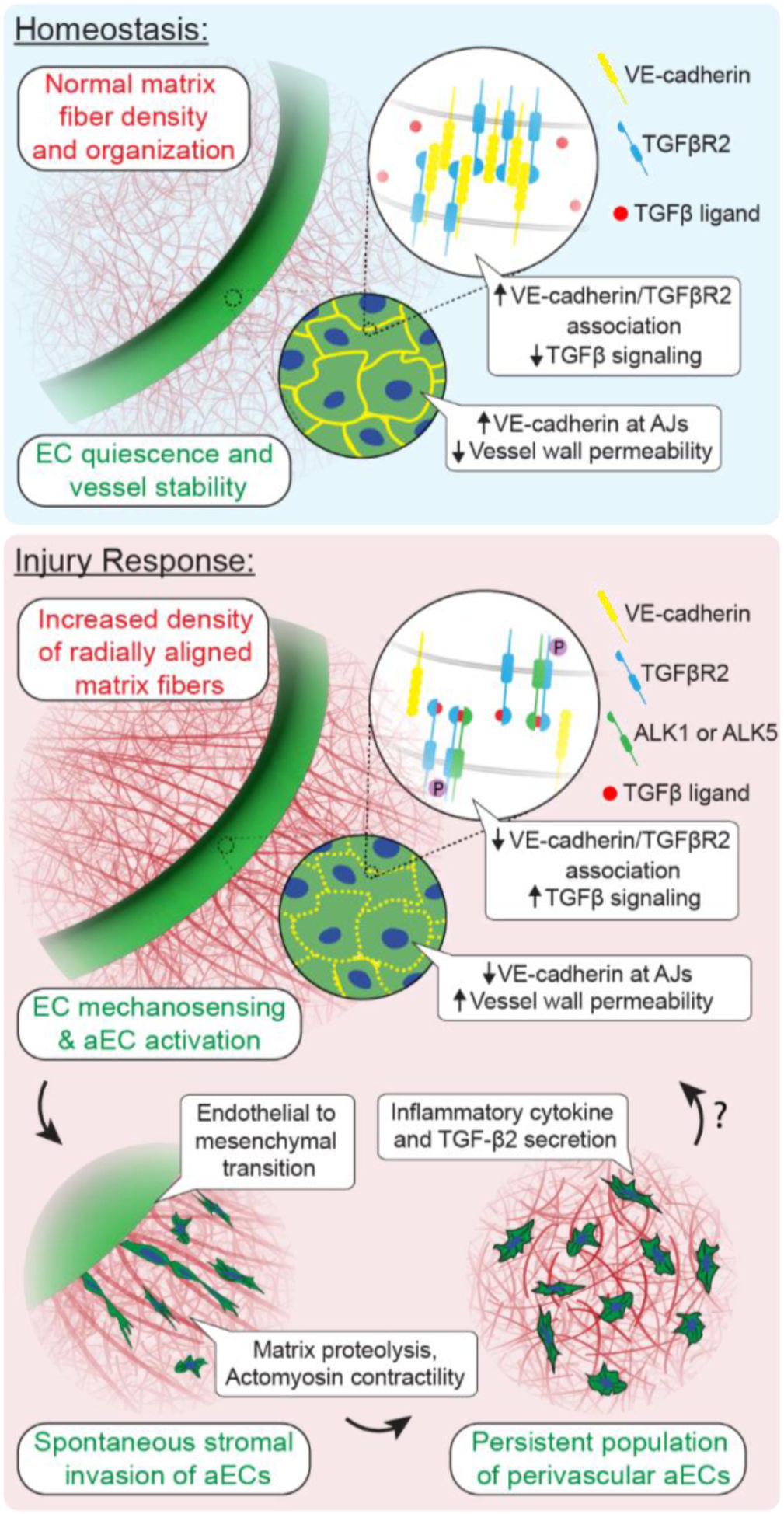
Summary schematic of key findings and outstanding questions (denoted with question mark).

A common observation across multiple animal models of lung fibrosis is microvasculature rarefaction towards the late-stage of the disease, during which both fiber density and TGF-β levels increase^74–76^. Our data demonstrates that ECs engaging a fibrous topography destabilize VE-cadherin which renders ECs more susceptible to TGF-β ligands and downstream signaling. The finding that VE-cadherin localization to AJs confers protective shielding from TGF-β-induced apoptosis may offer a promising therapeutic approach to prevent microvascular injury, rarefaction, and resulting tissue hypoxia in fibrotic diseases.

## DISCUSSION

While myriad soluble factors have been identified as activating signals for quiescent ECs and mediators of tip cell invasion ^77^, the role of physical cues presented by the ECM in directly driving this process or synergizing with soluble cues is far less established. EC activation and changes in ECM properties, such as fibrillar matrix density and organization, have both been observed following wound healing and during pathologic processes such as fibrosis. Here, we demonstrate how perivascular ECM fiber density directly influences the state of arteriole/venule-scale endothelium. Using an MPS platform of microvessels embedded within a tunable fibrous hydrogel composite, we examined the EC mechanosensing response to perivascular fiber density and found that heightened fiber density increased YAP-mediated mechanosensing, EndMT signaling, and the invasion of which failed to form lumenized capillaries and were often individualized cells, similar to those noted in in vivo lineage tracing studies. Mechanistically, elevated fiber density increased the destabilization of VE-cadherin containing AJs, diminished vessel barrier function, and increased EC TGF-β signaling via disruption of a VE-cadherin–TGF-βR2 interaction. Further ATECs within and engaging matrix with heightened fiber density possessed transcriptomic profiles suggesting a pro-inflammatory state and secreted TGF-β2. Altogether, our work identifies how enhanced fiber density associated with fibrogenesis regulates EC phenotype to generate ATECs which may play unappreciated roles in reinforcing fibrotic cascades.

Fibrotic tissues have been characterized by microvascular abnormalities, but the relationship between angiogenesis and fibrogenesis are still widely debated and little is known about the EC signaling that underlies these changes. Many studies report an increased angiogenic response based on increases in EC numbers (eg. CD31^+^ cells in a histological section) within a given injured or fibrotic tissue, but careful analysis of the connectivity, perfusability, permeability, and thus overall functional state of these newly formed EC structures has not been previously performed. Here, in a bleomycin-induced lung injury model, we show that Tie2Cre-mTmG lineage-traced ECs in the murine lung markedly expand at the peak of the bleomycin-induced fibrotic response, as marked by α-SMA^+^ MFs and matrix deposition. Coinciding with this expansion, we observed increased fibrillar ECM surrounding bronchial vasculature and the appearance of individualized, αSMA^+^ ATECs adjacent to bronchial arterioles and venules. Previous single cell RNA sequencing studies focused on interstitial lung disease patients have identified the intriguing expansion of ECs, which were broadly termed peribronchial ECs. Why ECs resident to large airway-associated vasculature are uniquely susceptible to fibrosis-associated activation remains unclear; however, one working model informed by our findings in the bleomycin and MPS models is a greater sensitivity to perivascular ECM remodeling in response to injury. Perhaps related to this, blood flow through the bronchial venous system is significantly (∼30%) reduced due to shunting of incoming blood flow to the pulmonary vasculature. Correspondingly, bronchial venules possess thin walls with limited adventitia – this lack of mural matrix support may increase EC sensitivity to changes in the perivascular matrix and/or better enable ATEC invasion into the perivascular space.

Through MPS modeling and transcriptomic analyses, we find that fibrous ECM drives the formation of ATECs via matrix mechanosensing and EndMT, which enables the proteolysis-dependent invasion of ATECs. In mice, EC activation by bleomycin-induced injury occurs through the activation of YAP/TAZ, and here we demonstrate the YAP inhibitor DMF blocks the formation of ATECs in fibrous hydrogels, presumably through inhibiting YAP-mediated mechanotransduction. Previous studies in lung and other organs suggest a role for EndMT in fibrosis by directly bolstering the myofibroblast population. In contrast, here we posit that EndMT mediated by fibrous ECM promotes the formation of a disease-mediating ATEC phenotype. Transcriptomic analysis, ELISA, and cytokine profiling suggest that fibrous topography transition ECs towards a source of TGF-β2 and other proinflammatory cytokines, consistent with single cell RNAseq studies in bleomycin-treated mouse lung identifying expanded EC populations derived from preexisting bronchial venous ECs with specialized inflammatory functions ^78^. It is therefore plausible that ATECs may play a key role in reinforcing fibrotic cascades through signaling interactions with the neighboring microvasculature and other stromal cells including fibroblasts and macrophages. Supporting an EC-centric feedback loop, simulating ATEC secretion of TGF-β2 with exogenous addition of this pro-fibrotic growth factor resulted in elevated apoptosis in parent microvessels surrounded by matrix possessing heightened fiber density. As TGF-β2 and fibrillar collagen density are both elevated during later phases of fibrosis^79^, these studies provide insight to the molecular mechanisms that may explain the common observation of microvasculature rarefaction.

As local changes in ECM in response to bleomycin and in human idiopathic lung fibrosis is highly heterogenous across the lung, we employed fiber-hydrogel composites within vascular MPS to better control the local microenvironment towards better understanding underlying pathophysiology. Using this approach, we found that adhesion of ECs within parental vessels to surrounding fibers led to decreased vessel barrier function and junctional weakening through increased turnover of VE-cadherin AJs. How EC adhesion to fibers as versus mechanical modulation of a surrounding bulk hydrogel distinctly modulates the EC response is an interesting and important mechanobiological question open for future investigation. Cell adhesion to ECM fibers in other contexts has recently been demonstrated to result in focal adhesions with distinct integrin and intracellular plaque compositions^80^, and this is consistent our observed differences for EC plated on flat or fibrous substrates sharing the same adhesive ligand. Simulating increased subendothelial stiffness using tailored 2D polyacrylamide hydrogels has been reported to increase EC TGF-β2 secretion ^81^, which we also observe to increase in response to heighten fiber density. Therefore, it is plausible that both conventional ECM mechanosensing pathways and to-be-characterized fiber-associated focal adhesion signaling cooperate to orchestrate EC AJ instability and downstream gene regulatory network changes underlying ATEC formation.

This work describes a novel VE-cadherin-gated mechanism controlling TGF-β signaling dependent on AJ stability. Using mass spectrometry and confirming with co-immunoprecipitation experiments, we found that TGF-βR2 is an abundant interacting protein with VE-cadherin. Intriguingly, this interaction is modulated by EC adhesive state, as evidenced by decreased co-immunoprecipitation of TGFβ-R2 with VE-cadherin in ECs engaging fibrous ECM, presumably due to the destabilization of VE-cadherin. In turn, EC adhesion to fibrous ECM led to destabilization of VE-cadherin-rich AJs and increased TGF-β signaling as evidenced by elevated TGF-βR2 phosphorylation and downstream SMAD3 nuclear localization. Whether VE-cadherin interactions with TGFβ-R2 block engagement of the receptor to TGF-β ligand or prevents TGF-βR2 receptor heterodimerization to suppress receptor signaling remains to be determined. However, our finding agrees with a recent study describing how Neuropilin-1 suppresses EC activation and TGF-βR2-dependent SMAD2/3 signaling through stabilization of VE-cadherin AJs^82^. This suggests that conditions that impair AJs and VE-cadherin stability may potentiate EC TGF-βR2-dependent TGF-β signaling, such as in the contexts of inflammatory cytokines histamine or thrombin and VEGF demonstrated here, which may be relevant to vascular inflammation and atherosclerosis^83^. Altogether, these studies provide new insights into how physical microenvironmental cues regulate EC phenotype, how pro-inflammatory ATECs may contribute to the pathogenesis of fibrosis, and identify new mechanisms that that may be targetable for treating pulmonary fibrosis as well as other fibrotic diseases.

## MATERIALS AND METHODS

### Reagents

All reagents were purchased from Sigma-Aldrich and used as received, unless otherwise stated.

### Lung injury model in a mouse line enabling EC lineage tracing

Male B6.Cg-Tg(Tek-cre)1Ywa/J transgenic mice were crossed with female Gt(ROSA)26Sortm4(ACTB-tdTomato,-EGFP)Luo mice to create Tie2cre/mTmG mice where all cells at baseline express membrane-localized tdTomato, but upon Tie2-driven Cre recombinase expression, flip expression to membrane-localized GFP. 8-12 week-old male mice were injected intratracheally with a single dose of 0.04 units bleomycin solubilized in PBS. Control mice lung were collected 3 weeks after dosing with PBS. Injured mice lungs and blood were harvested at week 3 after bleomycin instillation to examine the peak fibrotic response following lung injury. Warm PBS was perfused through the left ventricle at 5 ml/min until the liver turned white to clear lung tissue of red blood cells. Lungs were then inflated with 1.5ml of 2% w/v low melting agarose (Sigma A9025) in 4% w/v PFA via a catheter inserted into the upper trachea. Upon agarose gelation, the entire lung was extracted and immediately fixed in 4% w/v PFA overnight at 4°C then washed with PBS three-times the next day and stored in PBS with 0.01% sodium azide at 4 °C until PCLS sectioning.

### Tissue sectioning and advanced cubic clearing

The entire lung was first separated into left and right lobes then glued vertically on specimen holder to reflect original anatomical position. Lobes were then embedded in 2% w/v low melting agarose in PBS, to approximately match the stiffness of the tissue with the surrounding material during vibratome sectioning. PCLS (200 μm) were cleared using the advanced cubic clearing method^84^. Specifically, lung sections were cleared in dH2O with 50% CUBIC-L containing 10% w/w N-butyldiethanolamine, and 10% w/w Triton X-100 in dH2O for one day, and then 100% CUBIC-L for another day. Following clearing, lung tissue was permeabilized for one day in PBS solution containing 20% v/v DMSO, 0.1% v/v tween-20, 0.1% v/v triton X-100, 0.02% v/v SDS, 0.1% w/v deoxycholate and 0.1% w/v tergitol NP40. Tissue sections were blocked and stored in PBS solution containing 0.2% v/v TritonX-100, 6% v/v goat serum, and 10% v/v DMSO until staining.

### Lung tissue immunofluorescent staining and imaging

Tissue was incubated with anti-αSMA antibody (1:1000, Abcam #AB7817) or F4/80 antibody (1:250, Cytoskeleton #70076S) overnight in PBS containing 1 mg/ml heparin, 0.2% v/v tween-20, 5% v/v DMSO, 3% v/v goat serum, and 1% w/v bovine serum albumin followed by a 12h PBS wash overnight and overnight incubation in buffer containing Alexa-conjugated secondary antibodies (1:1000, Invitrogen #A-21133). Tissue was then stained by DAPI and NHS-ester (1:3000, Invitrogen# A20006) sequentially, each for 30 minutes at room temperature with 3x PBS washes in between. Tissues were imaged in CUBIC-R+M composed of 45% w/w antipyrine, 30% w/w N-methylnicotinamide, and 0.5% v/v N-butyldiethanolamine in dH2O via a laser-scanning confocal microscope (Zeiss LSM800). TUNEL staining (Fisher, #C10619) following the manufacturer’s instructions was utilized for visualizing DNA damage. Lung tissue stained by TUNEL did not go through CUBIC clearing due to chemical incompatibility and 25 μm Z-stacks with 1 μm step size were taken using confocal at 20x only at the center of the tissue to avoid false positive signals at the plane of sectioning.

### Organotypic model device fabrication

Device moulds were designed in AutoCAD and printed via stereolithography (SLA) by Protolabs (Maple Plain, MN). Polydimethylsiloxane (PDMS, 1:10 crosslinker:base ratio) devices were replica-casted from SLA printed moulds, cleaned with isopropyl alcohol and ethanol, and bonded to glass coverslips with a plasma etcher. Devices were treated with 0.01% (w/v) poly-l-lysine and 0.5% (w/v) L-glutaraldehyde sequentially for 1 hour each to covalently crosslink ECM to the walls of the device, thus preventing potential hydrogel compaction from cell-generated forces. 160 µm stainless steel acupuncture needles (Lhasa OMS, Weymouth, MA) were dip-coated with 1% w/v gelatin to facilitate removal without hydrogel fracture, inserted into each device, and sterilized by UV ozone. Hydrogel precursor solution was then injected into each device and polymerized around needles. Hydrogels were hydrated in EGM2 media at 37℃ overnight (or greater than 12 hours) to dissolve the gelatin layer and needles were extracted to form 3D hollow channels fully embedded within a crosslinked hydrogel and positioned 500 µm away from PDMS and glass boundaries. A chilled solution of 100 µg/ml matrigel diluted in PBS was perfused through channels and allowed to adsorb onto the hydrogel channel surface at 4°C overnight. Residual Matrigel was then rinsed with PBS twice.

### Dextran vinyl sulfone polymer synthesis

Dextran (MW 86,000 Da, MP Biomedicals, Santa Ana, CA) was functionalized with vinyl sulfone groups as in ^45,46,85^. Briefly, dextran (5 g) was dissolved in 0.1 M sodium hydroxide solution (250 mL) at room temperature. Divinyl sulfone (3.875 ml, ThermoFisher Scientific, Waltham, MA) was added and the reaction was carried out for 4 minutes with vigorous stirring (1500 RPM) at room temperature. The reaction was terminated by adjusting the pH to 5.0 with the addition of hydrochloric acid. The reaction product was dialyzed against milli-Q water for 3 days with two water exchanges daily. The dialyzed reaction product was lyophilized for 3 days and characterized by ^1^H-nuclear magnetic resonance spectroscopy in D_2_O. The degree of vinyl sulfone functionalization was calculated as the ratio of the proton integral (6.91 ppm) and the anomeric proton of the glucopyranosyl ring (5.166 and 4.923 ppm); a vinyl sulfone/dextran repeat unit ratio of 0.66 was determined.

### Fiber segment fabrication

DexVS fiber segments were generated as in Matera et al ^45^. DexVS was dissolved at 0.6 g ml^−1^ in a 1:1 mixture of Milli-Q water and dimethylformamide with 0.015% Irgacure 2959 photoinitiator and 0.5 mM methacrylated rhodamine to enable visualization (Polysciences Inc., Warrington, PA). This polymer solution electrospun into fibers within an environment-controlled glovebox held at 21°C and 30% relative humidity. Electrospinning was performed at a flow rate of 0.3 ml hour^−1^, gap distance of 5 cm, and voltage of −10.0 kV onto a grounded collecting surface attached to a linear actuator. Fiber mats were collected on glass slabs and primary cross-linked under ultraviolet light (100 mW cm^−2^). After polymerization, fiber segments were resuspended in a 3 ml volume of PBS. The total volume of fibers was then calculated via a conservation of volume equation: total resulting solution volume = volume of fibers + volume of PBS. After calculating total fiber volume, solutions were re-centrifuged, supernatant was removed, and fiber pellets were resuspended to create a 10% v/v fiber stock solution, which were then aliquoted and stored at 4°C. To enable cell adhesion, 2 mM arginylglycylaspartic acid (RGD, CGRGDS; GenScript, George Town, KY) was coupled to vinyl sulfone groups along the DexVS backbone via Michael-type addition chemistry for 30 min, followed by quenching of excess VS groups in a 300 mM cysteine solution for 30 min.

### Fibrin-fiber composite hydrogels

Fibrin hydrogels were prepared with fibrinogen from bovine plasma dissolved in PBS at 50 mg ml^−1^ stock concentrations. 10 mg ml^−1^ fibrinogen was prepared in PBS and crosslinked with thrombin (6 units per mg of fibrinogen) for 20 minutes at 37℃. For fibrous hydrogels, DexVS fiber segments were incorporated within the fibrin hydrogel precursor solution over a 0-2% v/v density. To generate fiber alignment perpendicular to the long axis of the parent vessel (0° alignment), fibrin precursor solution containing fibers were injected such that flow across acupuncture needles (rigid cylinder) generated a flow profile that aligned fibrils in the direction of flow. To generate fiber alignment parallel to the long axis of the parent vessel (90° alignment), fibrin precursor solution containing fibers were first injected into the device. Subsequent insertion of acupuncture needles generated flow profiles to align fibers in the direction of needle insertion. To generate collagen bundle-embedded fibrin hydrogels, type I collagen rat tail (4.8 mg ml^−1^) was adjusted to a pH of 7.4 with sodium hydroxide and PBS on ice. Next, 80°C MQ water was added in a 1:1 ratio (2.4 mg ml^−1^ final concentration) and immediately vortexed for 30 seconds. Collagen bundles formed immediately during vortexing and were isolated via centrifugation (500 g for 4 minutes).

### Mechanical testing

The Young’s modulus of composite hydrogels were measured by atomic force microscopy (AFM; Nanosurf, Liestal, Switzerland) in contact mode. Indentations were made at a loading rate of 2 µm/s with silicon nitride cantilevers (AppNano, Mountain View, CA) with a nominal spring constant of 0.046 N/m and a 5 μm diameter spherical glass bead. Force-displacement curves were taken at a minimum of 3 regions on each hydrogel and fit to the Hertz model assuming a Poisson’s ratio of 0.5 to estimate the elastic modulus.

### Device seeding and culture

Human umbilical vein, liver, lung, and dermal endothelial cells (Lonza, Switzerland) were cultured in endothelial growth media (EGM2, Lonza). HUVECs were passaged upon achieving confluency at a 1:4 ratio and used in studies from passages 4 to 9. Liver, lung, and dermal ECs were passaged at a 1:4 ratio and used in studies from passages 2 to 6. A 10 µl solution of suspended ECs (10 million cells ml^−1^ density) was added to one reservoir of the endothelial channel and inverted for 30 minutes to allow cell attachment to the top half of the channel, followed by a second round of seeding with the device upright for 30 minutes to allow cell attachment to the bottom half of the channel. ECs reached confluency and self-assembled into stable parent vessels over 24 hours. Media and chemokines were refreshed every 24 hours and devices were cultured with continual reciprocating flow utilizing gravity-driven flow on a seesaw rocker plate at 0.33 Hz. Tip cell formation in response to S1P gradients was assessed by adding 0-50 nM S1P diluted in EGM2 media added only to the chemokine channel for 3 days beginning 16 hours after cell seeding. Fiber-induced tip cell formation studies were cultured in EGM2 media alone for 3 days after 16 hours post cell seeding. Agents for permeability agonist and pharmacologic dosing studies were incorporated into EGM2 media and added to both the endothelial and chemokine channels for 3 days after 16 hours post cell seeding. For studies examining the effect of supporting mural cells on EC activation, human mesenchymal stem cells (hMSCs) were cultured in high-glucose DMEM containing 5% fetal bovine serum and 1% P/S and were passaged upon achieving confluency at a 1:3 ratio and used from passages 3-6. hMSCs were first seeded into device channels as described above, at a 10 µl solution of 1.5 million cells ml^−1^. Endothelial cells were seeded as described above 1 hour after hMSCs seeding and attachment.

### Diffusive permeability measurements

Diffusive permeability was quantified as in Polacheck et al ^86,87^. Briefly, fluorescent dextran (70 kDa Texas Red, Thermo Fisher) was dissolved in EGM2 media at 12.5 µg ml^−1^, perfused through endothelialized channels, and imaged at 1 second intervals to measure the flux of dextran across the endothelium into the ECM. The resulting diffusion profile was fit to a dynamic mass-conservation equation as in ^88^ with the diffusive-permeability coefficient (*P*_D_) defined by *J* = *P*_D_(*c*_vessel_ − *c*_ECM_), where *J* is the mass flux of dextran, *c*_vessel_ is the concentration of dextran in the vessel, and *c*_ECM_ is the concentration of dextran in the perivascular space.

### Western blotting

Samples for western blotting were collected from confluent EC monolayers cultured on tissue culture plastic or electrospun fiber matrices^89^ for 2 days after cell seeding and were treated with either EGM2 media, or EGM2 media containing 50 ng ml^−1^ VEGF (Peprotech, Cranbury, NJ), 50 ng ml^−1^ TNFα, or 2 U ml^−1^ Thrombin. To collect protein lysates, cells were collected with a cell scraper in chilled PBS solution and centrifuged to generate cell pellet. Supernatant was removed and replaced with RIPA buffer containing protease and phosphatase inhibitors. Following freeze fracture of cell membranes (10 minutes at –80°C), samples were centrifuged for 10 minutes at 20,000 g, and the supernatant was collected which contained the purified protein lysate. Protein lysate concentrations were measured using a BCA assay, and 30 µg ml^−1^ protein was loaded into a 4-20% Tris-Glycine Novex wedge well. Proteins were separated with electrophoresis for 90 minutes at 120V in running buffer. Protein gels were then transferred to a PVDF membrane for 60 minutes at 20V in transfer buffer at 4°C. PVDF membranes were then blocked in 5% w/v blocking grade milk buffer, and stained with an anti-VE-cadherin primary antibody (sc-9989, 1:1,000, Santa Cruz Biotechnology, Dallas, TX) and β-tubulin (66240-1-Ig, 1:10,000, Proteintech, Rosemont, IL) overnight on an orbital shaker at 4°C. Membranes were then rinsed in TBST and stained with HRP and imaged on a gel imager.

### Transcriptomic analysis

Samples for microarray analysis included 1) confluent EC monolayers cultured on 10 mg/ml fibrin hydrogel slabs, 2) confluent EC monolayers cultured in channels within 10 mg/ml fibrin hydrogels, 3) encapsulated, single ECs embedded within FD 0% 10 mg ml^−1^ fibrin hydrogels, and 4) encapsulated, single ECs embedded within FD 2% 10 mg ml^−1^ fibrin hydrogels. All conditions were cultured for 3 days upon which nattokinase (100 fibrinolytic units ml^−1^) was incorporated into EGM2 media for 15 minutes to digest the surrounding fibrin hydrogel to release attached ECs. ECs were pelleted and RNA isolation was performed via RNeasy mini kit per manufacturer’s protocol. Purified RNA samples were submitted to the University of Michigan Sequencing Core for microarray analysis on an Affymetrix chip. Gene expression data was analyzed utilizing Advaita Bioinformatics software^90^.

### Cytokine and growth factor secretion analysis

To assess how fibrous topography influenced EC cytokine secretion, we employed a recently established microfluidic ELISA and an antibody cytokine detection array membrane. For both assays, samples were cultured for 7 days, with supernatant media refreshed every 2 days. Thus, conditioned media collected for analysis contained 2 days of cell secreted factors (i.e. days 5 –7). As cell-secreted factors, such as TGF-β2 become bound to the extracellular matrix (i.e. latent TGF-β complexes) upon secretion, the fibrin hydrogel was digested utilizing nattokinase as described above. To stabilize TGF-β2 cytokines, hydrochloric acid was added to conditioned media to pH 5 only for TGF-β2 measurements. All samples were stored at –80°C and thawed immediately prior to cytokine detection assays. Microfluidic-based ELISA was performed as previously described in detail in Tan et al ^91^. Inflammatory cytokine detection array membrane was performed per manufacturer protocol (R&D Systems, Proteome Profiler Human Cytokine Array, #ARY005B).

### VE-cadherin/TGF-β2 signaling studies

For studies examining SMAD3 localization, ECs were first treated with mitomycin C (20 µg ml^−1^ for 2 hours) to inhibit cell proliferation and thus control over cell seeding densities during culture. For variable cell density SMAD3 studies, ECs were seeded onto 2D 10 mg ml^−1^ fibrin hydrogels (18 mm coverslips treated with poly-l-lysine and l-glutaraldehyde) at a low (25 K cells cm^−2^), medium (50 K cells cm^−2^), or high (100 K cells cm^−2^) cell density. The following day, samples were treated with 0 or 10 ng ml^−1^ TGF-β2 (Peprotech) in EGM2 media (without serum or fibroblast growth factor (FGF), as FGF has previously been shown to inhibit TGF-β signaling) for two days with media refreshed daily. For mosaic SMAD3 studies, scrambled control and VECKO ECs were seeded at varying ratios (0, 50, 75, and 100% VECKOs) at a constant 100 K cells cm ^−2^ density onto 2D 10 mg ml^−1^ fibrin hydrogels. For fibrin hydrogel vs fibrin hydrogel + fibers studies, fibrin hydrogel + fibers conditions were generated by electrospinning DexVS fiber matrices onto a dried gelatin-coated coverslip. A solution of fibrin hydrogel was then added to the DexVS fiber matrix with a second glutaraldehyde treated coverslip placed on top. After fibrin polymerization and hydration, the dried-gelatin coverslip was removed, resulting in a DexVS fiber matrix adhered on top of a fibrin hydrogel-coated glutaraldehyde coverslip. ECs were seeded on to fibrin or fibrin + FD 2% hydrogels at a 100 K cells cm^−2^ density and dosed with TGF-β2 identical as described above. For scratch wound studies, glass coverslips were seeded with ECs at 100 K cells cm^−2^ density and scratched with a P1000 tip the following day and dosed with TGF-β2 identical as described above except only for 1 day as at longer timepoints ECs were observed to close the scratch region.

### Fluorescent staining

Samples were fixed with 4% paraformaldehyde and permeabilized with a PBS solution containing Triton X-100 (5% v/v), sucrose (10% w/v), and magnesium chloride (0.6% w/v) for 1 hour each at room temperature. AlexaFluor 488 phalloidin (Life Technologies, Carlsbad, CA) was utilized to visualize F-actin. 4’, 6-diamidino-2-phenylindole (DAPI, 1 µg ml^−1^) was utilized to visualize cell nucleus. To visualize VE-cadherin, YAP, SNAI1, vimentin, or SMAD3, samples were sequentially blocked in bovine serum albumin (0.3% w/v), incubated with primary antibody [mouse monoclonal anti-VE-cadherin (1:500, Santa Cruz Biotechnology), mouse monoclonal anti-YAP (1:500, Santa Cruz Biotechnology), rabbit monoclonal anti-SNAI1 (1:500, Cell Signaling Technologies), mouse monoclonal anti-vimentin (1:500) rabbit monoclonal anti-SMAD3 (1:500, Cell Signaling Technologies)], and incubated with secondary AlexaFluor 647 goat anti-mouse or anti-rabbit IgG (H+L) (1:1000, Life Technologies) each for 1 hour at room temperature. For VE-cadherin pulse studies, a mouse monoclonal anti-VE-cadherin antibody (55-7H1 clone) pre-conjugated with AlexaFluor 647 was added to cell culture media (1:500) for 30 minutes followed by media rinses as described in ^92^.

### Microscopy and image analysis

Fluorescent images were captured on a Zeiss LSM800 confocal microscope. To quantify tip cell formation, the number and distance of tip cells were measured in FIJI. Tip cells were defined morphologically as the leading cell of an invading strand or as a single invading cell. VE-cadherin signal intensity was quantified by summing the total VE-cadherin signal and normalizing to the number of cells in each field of view. Nuclear SMAD3 intensity was quantified by first masking the SMAD3 signal with a nuclear mask, then summing SMAD3 intensity and normalizing to the number of cells in each field of view. Performing these analyses on a field of view basis allowed for Pearson’s correlation analysis between nuclear SMAD3 and VE-cadherin. Measurements of VE-cadherin junctional width was performed as described previously ^65,67,93^. Briefly, high resolution images of VE-cadherin were acquired on a confocal microscope. Images across conditions were thresholded under the same parameters. A line orthogonal to the long axis of the junction was drawn to measure the intensity profile and obtain VE-cadherin junction width.

### Statistics

Statistical significance was determined by one-way analysis of variance (ANOVA) or two-sided student’s t-test where appropriate, with significance indicated by p<0.05. Sample size is indicated within corresponding figure legends and all data are presented as mean ± standard deviation.

### Data availability

The data that support the findings of this study are available from the corresponding author upon reasonable request.

## AUTHOR CONTRIBUTIONS

WYW, JX, KAJ, MLK, and BMB designed all of the experiments. WYW, KAJ, EHJ, and DL conducted *in vitro* experiments and data analysis. JX, RNK III, EHS, AS, and BMB carried out in vivo studies and data analysis KL and CP provided support for imaging of western blots. HLH and CDD performed mechanical testing on hydrogels. XT and XF performed microfluidic ELISA measurements. KAJ and MLK performed VE-cadherin BioID, VE-cadherin CRISPR, TGF-βR2 experiments. WYW, JX, MLK, and BMB wrote the manuscript. All authors reviewed and edited the manuscript.

## ACKNOWLEDGMENTS

This work was supported in part by the National Institutes of Health (HL124322) and (GM150987). W.Y.W acknowledges financial support from the University of Michigan Rackham Merit Fellowship and National Science Foundation Graduate Research Fellowship Program (DGE1256260). C.D.D. acknowledges financial support from the National Science Foundation Graduate Research Fellowship Program (DGE1256260) and the Ruth L. Kirschstein National Research Service Award (F31HL152501).

