## supplementary data for "Perivascular matrix densification dysregulates angiogenesis and activates pro-inflammatory endothelial cells"

### SUPPLEMENTARY MATERIAL

The Supplementary Material includes 10 Supplementary Figures.

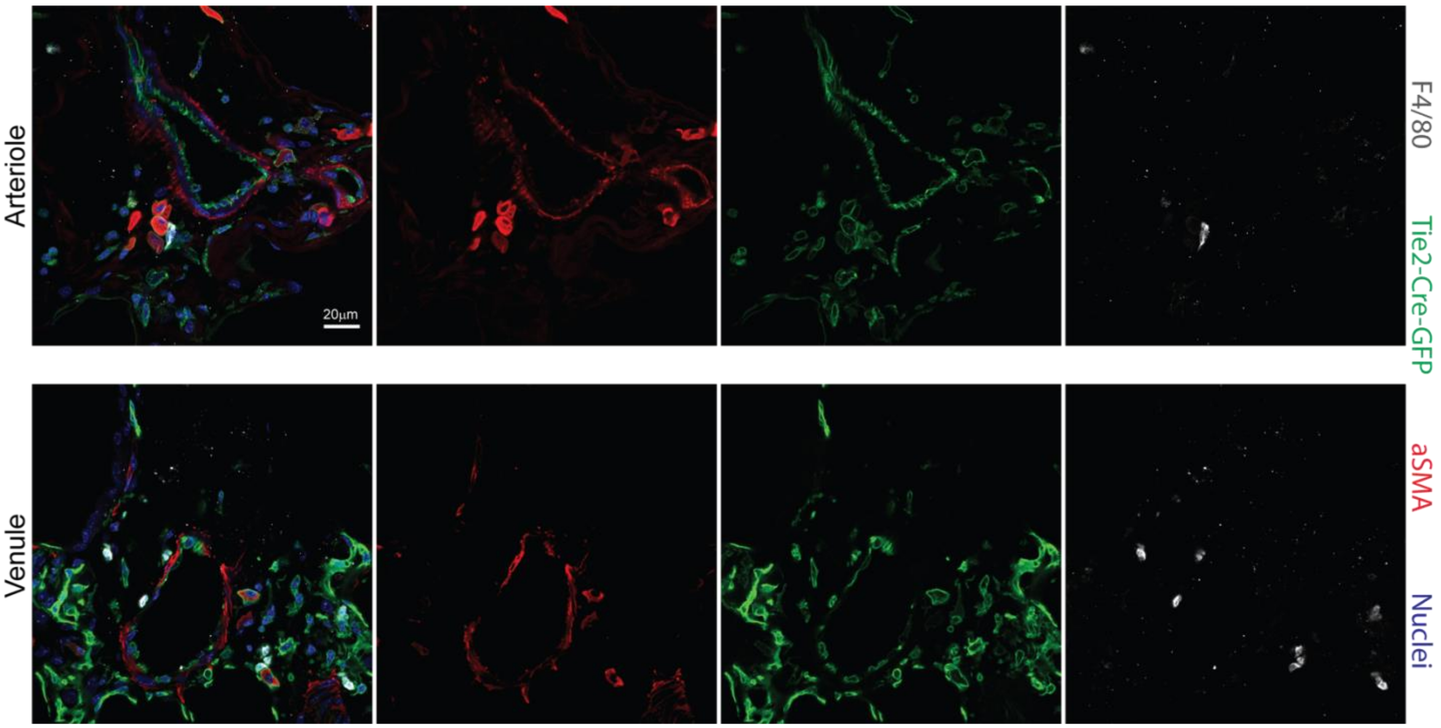


**Supplementary Figure 1: GFP/αSMA dual positive ECs are a distinct population from F4/80+ macrophage. a)** Representative images of an arteriole (top row) and venule (bottom row) with their perivascular spaces from week2 post bleomycin mice. 3D slicing views is included in supplementary videos. scale bar is 20um.

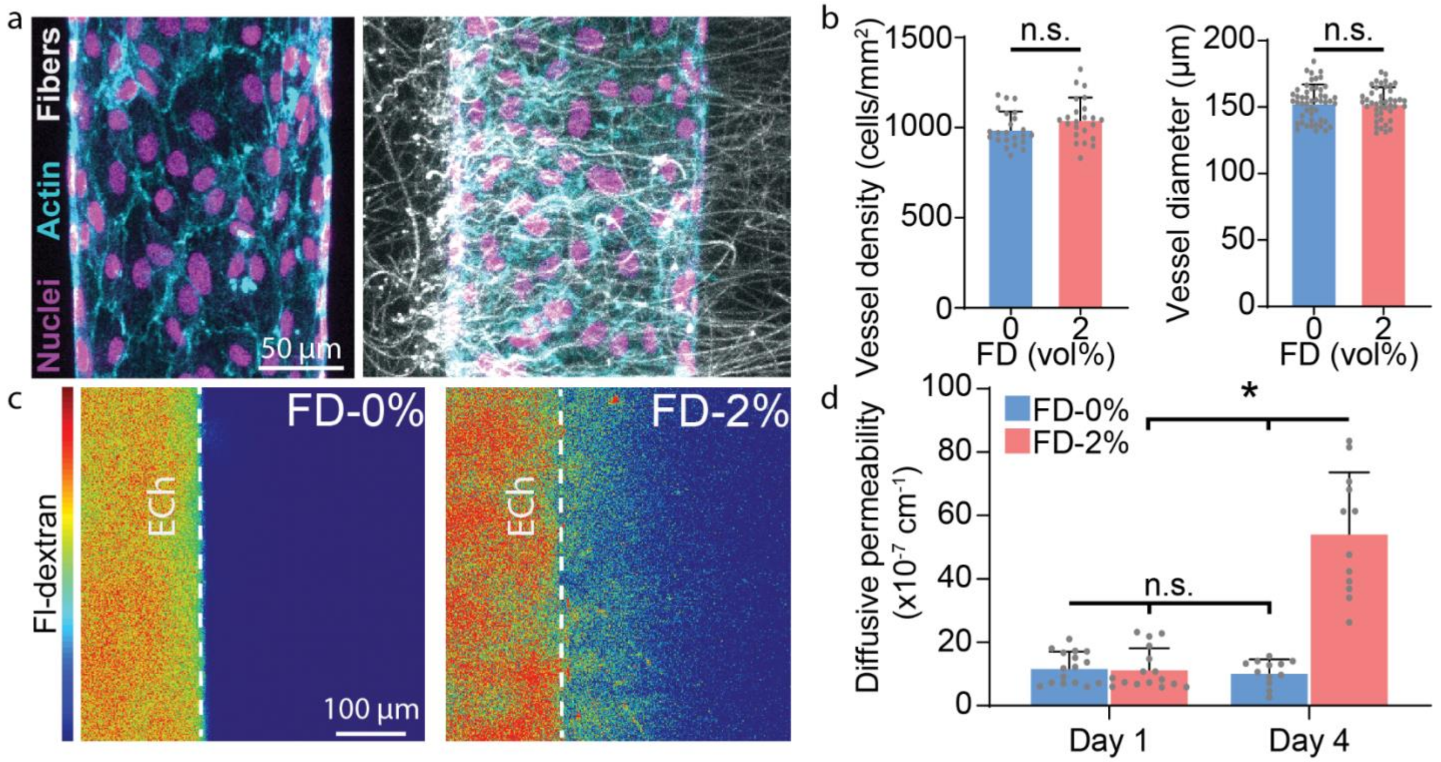


**Supplementary Figure 2: Analysis of microvessels in control vs. fibrous ECM after 16 hours of cell seeding. a)** Representative images (max intensity projection) of parent vessels cultured in control vs FD 2% hydrogels with channels coated in basement membrane proteins. **b)** Quantification of vessel cell density and diameter. **c)** Fluorescent dextran diffusion in control vs FD 2% hydrogels on Day 4 culture. **d)** Quantification of diffusive permeability of control and FD 2% hydrogels on Day 1 and 4. All data presented as mean ± std.; * indicates a statistically significant comparison with P<0.05. n.s. indicates a non-statistically significant comparison (b: two-sided student's t-test and d: one-way analysis of variance).

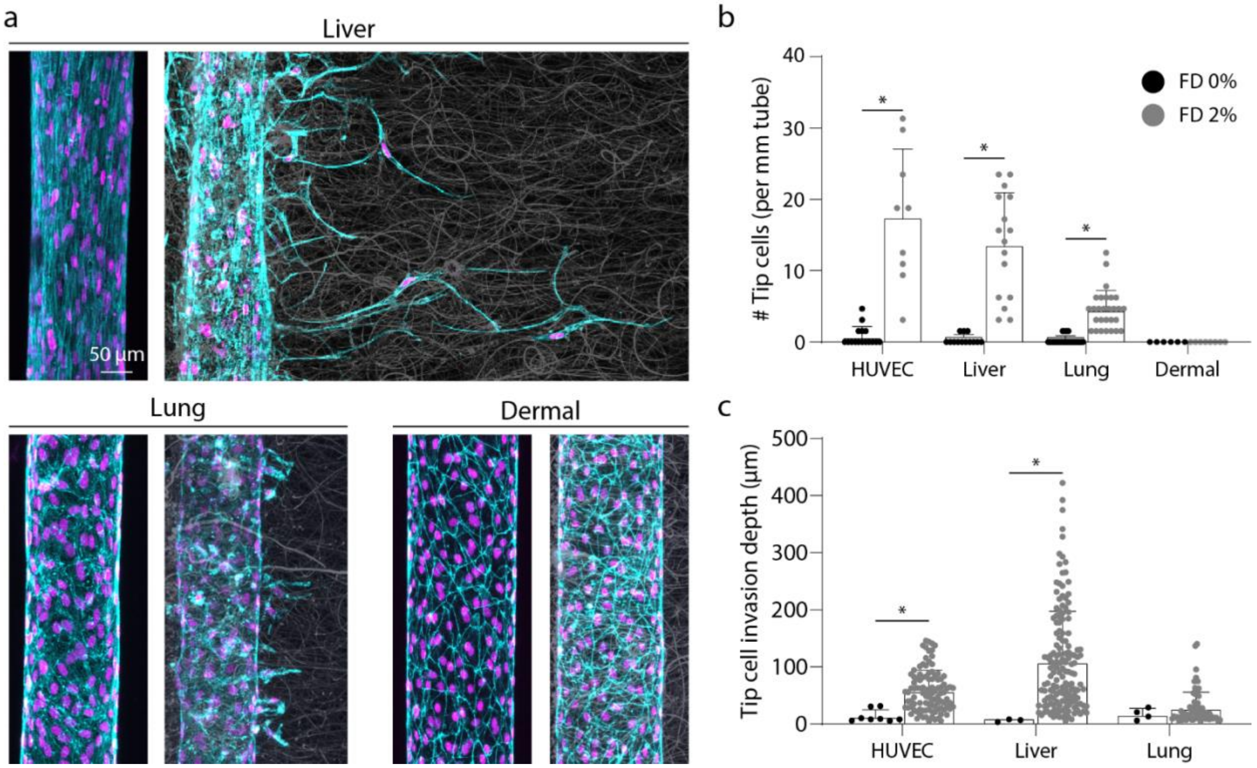


**Supplementary Figure 3: Fiber density promotes tip cell formation in liver and lung, but not dermal endothelial cells. a)** Representative images (max intensity projection) of liver, lung, and dermal endothelial cells cultured over 4-days in control or FD 2% hydrogels. Nuclei (magenta), F-actin (cyan). **b-c)** Quantification of number of tip cells and tip cell invasion depth for conditions in (a). All data presented as mean ± std.; * indicates a statistically significant comparison with $P<0.05$ (two-sided student's t-test).

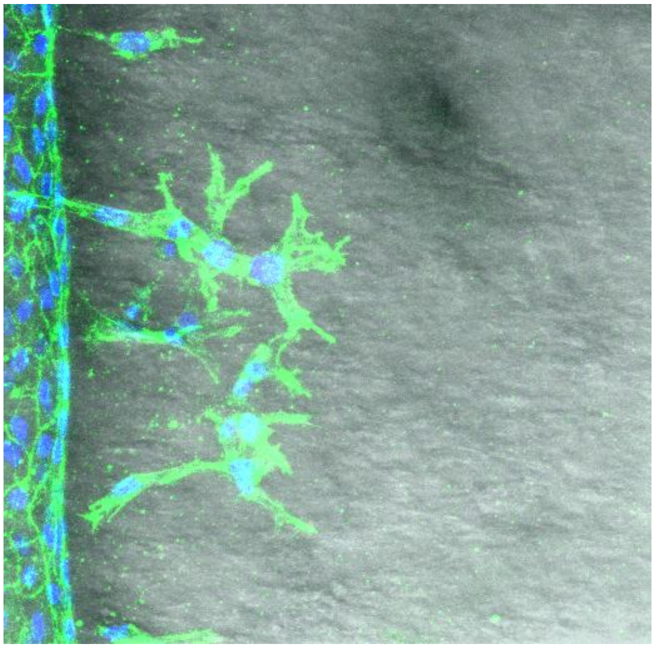

**Supplementary Figure 4: a)** Micrometer-scale **c**ollagen bundles were generated as in (Gong et al., 2020), isolated with centrifugation and embedded within 10 mg/ml fibrin hydrogels. Endothelial cells were cultured for 6 days. Nuclei (blue), F-actin (green), transmitted light (grey).

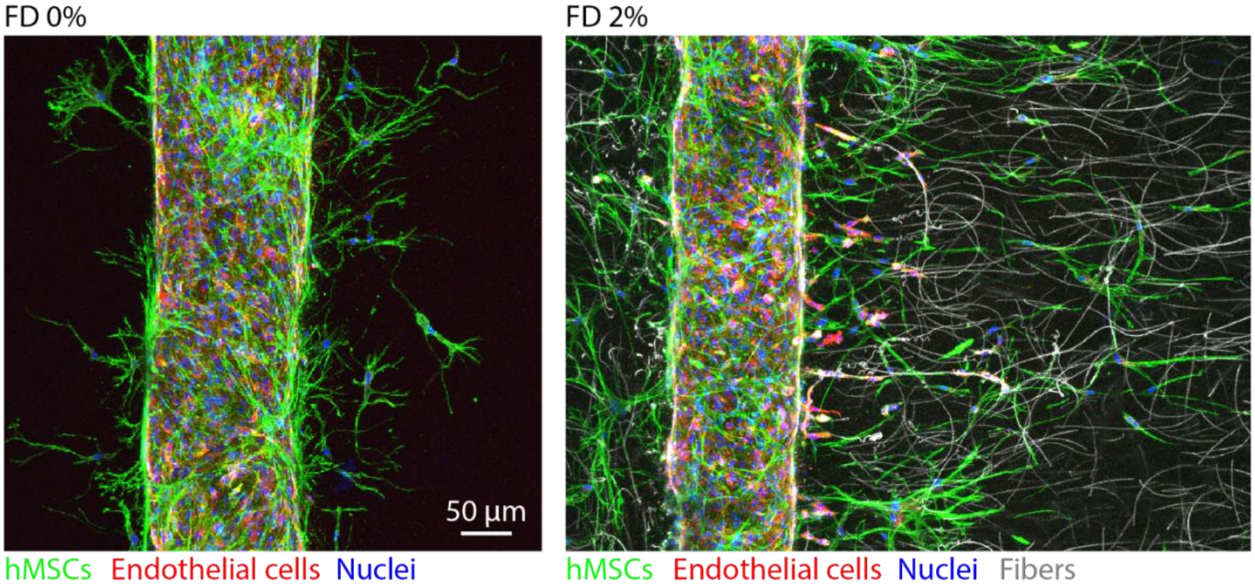


**Supplementary Figure 5: Fiber density promote tip cell formation with hMSC-coated endothelium.** Representative images (max intensity projection) of hMSCs and endothelial cells cultured over 4-days in control or FD 2% hydrogels.

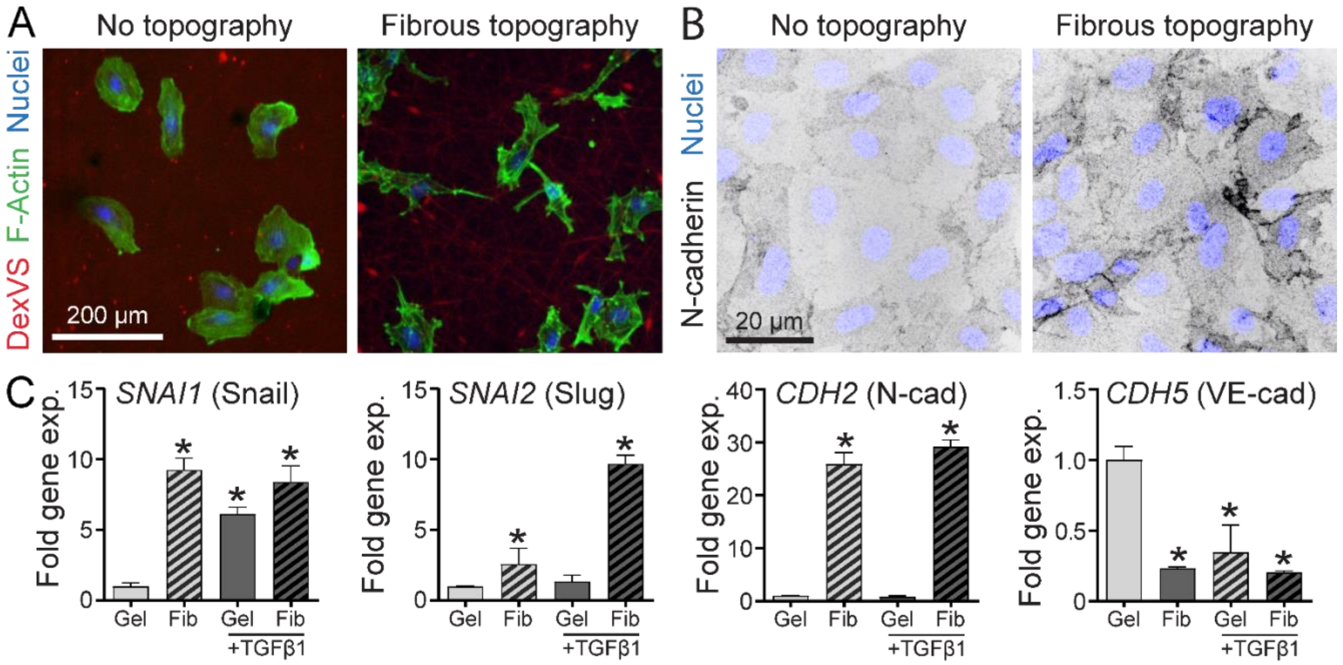


**Supplementary Figure 6: A)** Representative images (max intensity projection) of ECs seeded sparsely on flat DexVS hydrogels lacking topography and electrospun DexVS fibers, both functionalized with RGD to enable cell adhesion. B) Immunostaining of N-cadherin of ECs on identical substrates as in A. Note: intensity scale for N-cadherin in grayscale is inverted. C) Relative gene expression of sparsely plated ECs seeded on identical substrates as in A with or without exogenous TGF-β (5 ng/ml). Gene expression values normalized to flat DexVS gel substrates without TGF-β1 treatment.

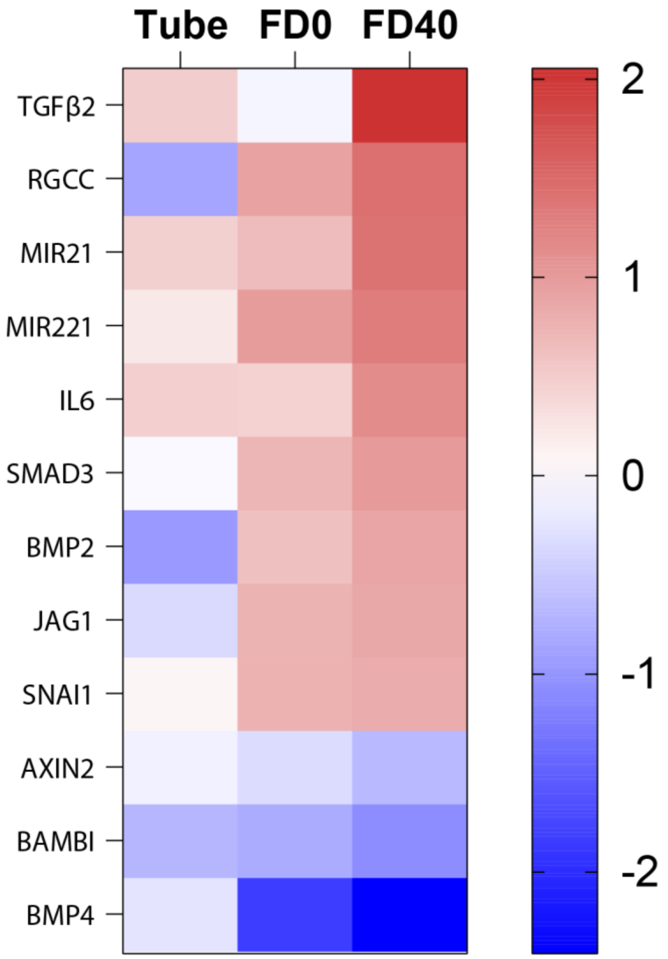


**Supplementary Figure 7:** Differentially expressed genes under "positive regulation of epithelial to mesenchymal transition" Gene Ontology category.

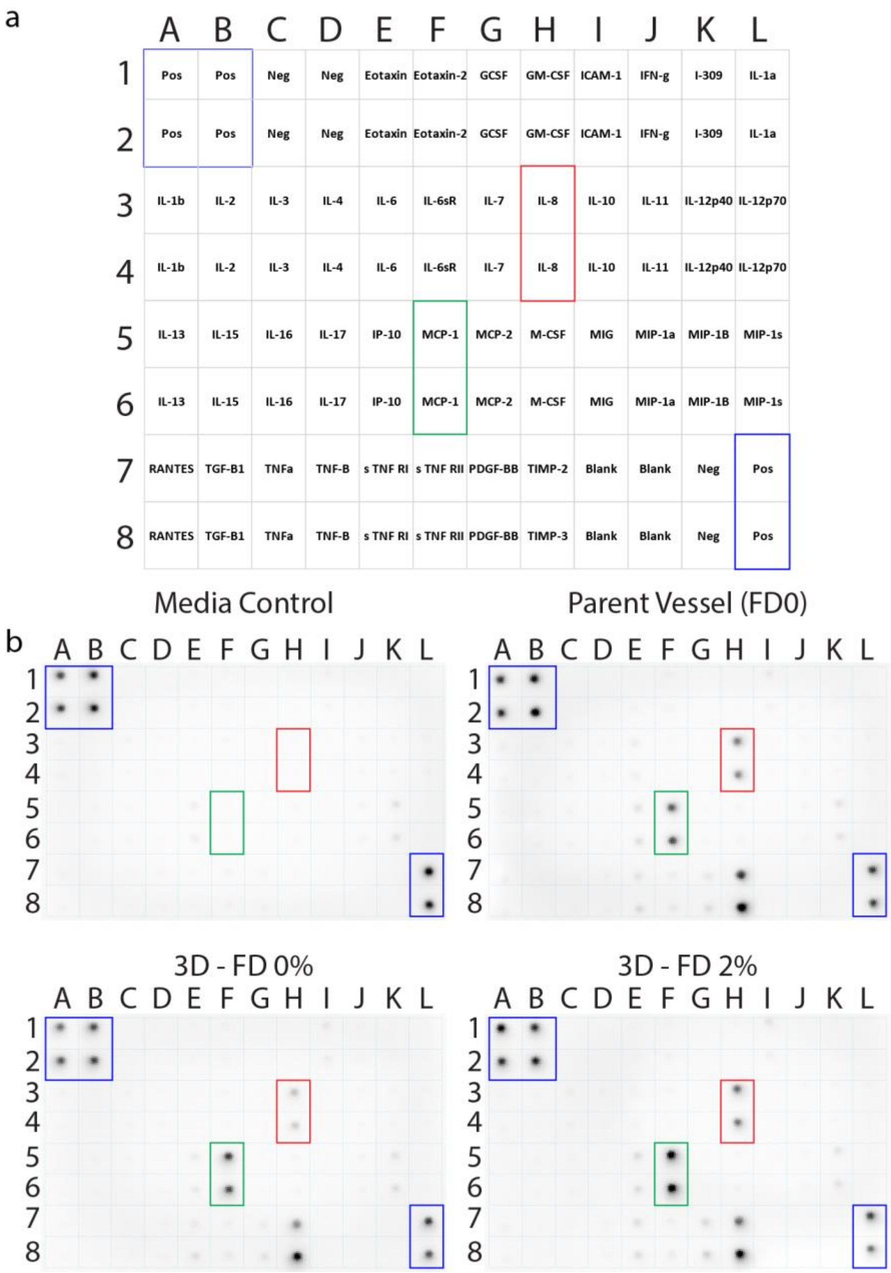


**Supplementary Figure 8: Human inflammation antibody membrane array. a)** Membrane array cytokine map. **b)** Intensity-based response of captured antibodies from media control, or conditioned media collected from cells cultured as a parent vessel, 3D embedded within control or fibrous hydrogels.

| Prey Protein | Normalized Total Spectra VE-cadherin-BI | Cytoplasmic BI |
|---|---|---|
| CTNNA1 | 194.61 | 3.1434 |
| JUP | 118.20 | 3.1434 |
| AFDN | 109.84 | 18.074 |
| **CDH5** | **100.29** | **0** |
| CTNNB1 | 87.155 | 0 |
| CTNND1 | 65.665 | 5.5009 |
| ERBIN | 37.011 | 3.1434 |
| MYO1C | 34.623 | 5.5009 |
| JCAD | 21.49 | 0 |
| ARHGAP21 | 20.296 | 3.1434 |
| SEPT9 | 20.296 | 4.751 |
| CKAP4 | 17.909 | 8.6443 |
| **TGFBR2** | **16.715** | **0** |
| LPP | 15.521 | 3.9292 |
| DLG5 | 14.327 | 4.7151 |

**Supplementary Figure 9: Top 15 VE-cadherin interactors identified via BioID and mass spectrometry anaylsis.**

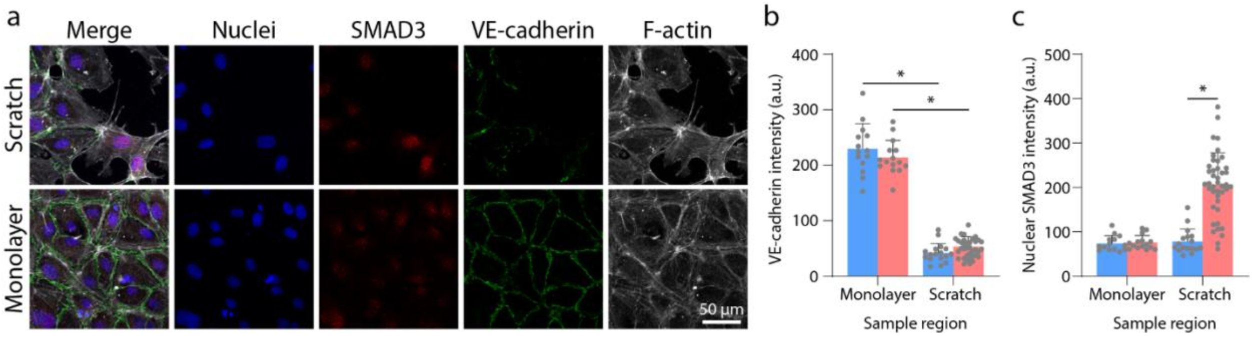


**Supplementary Figure 10: Scratch wound assay to modulate VE-cadherin stability. a)** Monolayer vs scratch regions within the same sample treated with 10 ng/ml TGFβ2 for 1 day. Nuclei (blue), SMAD3 (red), VE-cadherin (green), F-actin (white). **b-c)** Quantification of VE-cadherin and nuclear SMAD3 expression in monolayer vs scratch regions treated with or without 10 ng/ml TGFβ2. All data presented as mean ± std.; * indicates a statistically significant comparison with P<0.05 (one-way analysis of variance).
